# GVB+ neurons are resilient to tau-induced protein synthesis impairment

**DOI:** 10.1101/2025.03.28.645932

**Authors:** Jasper F.M. Smits, Thijmen W. Ligthart, Marta Jorge-Oliva, Skip Middelhoff, Fleur Schipper, Débora Pita-Illobre, Ka Wan Li, Wiep Scheper

**Affiliations:** Dept. of Human Genetics, Amsterdam UMC - Vrije Universiteit Amsterdam, The Netherlands; Dept. of Functional Genomics; Dept. of Molecular and Cellular Neuroscience, Center for Neurogenomics and Cognitive Research, Amsterdam Neuroscience, Vrije Universiteit Amsterdam, The Netherlands

**Keywords:** granulovacuolar degeneration bodies (GVBs), tau pathology, Casein kinase 1δ, proteostasis

## Abstract

In Alzheimer’s disease, many surviving neurons with tau pathology contain granulovacuolar degeneration bodies (GVBs), neuron-specific lysosomal structures induced by pathological tau assemblies. This could indicate a neuroprotective role for GVBs, however, the mechanism of GVB formation and its functional implications are elusive. Here, we demonstrate that CK1δ activity is required for GVB formation. CK1δ is sequestered in the GVB during this process in an autophagy-dependent manner. We show that neurons with GVBs (GVB+) are resilient to tau-induced impairment of global protein synthesis and are protected against tau-mediated neurodegeneration. GVB+ neurons do not exhibit differential activation of proteostatic stress responses, but have increased ribosomal content. Importantly, unlike neurons without GVBs, GVB+ neurons fully retain the capacity to induce long-term potentiation-induced protein synthesis in the presence of tau pathology. Our results have identified CK1δ as a key regulator of GVB formation that confers a protective neuron-specific proteostatic stress response to tau pathology. These findings provide novel opportunities for targeting neuronal resilience in tauopathies.

**Highlights:**

- GVB+ neurons are protected against tau-induced neurodegeneration
- CK1δ activity is essential for the formation of GVBs
- GVB+ neurons are resilient to tau-induced impairment of protein synthesis
- GVB+ neurons retain the capacity for LTP-regulated protein synthesis

**Graphical abstract:** 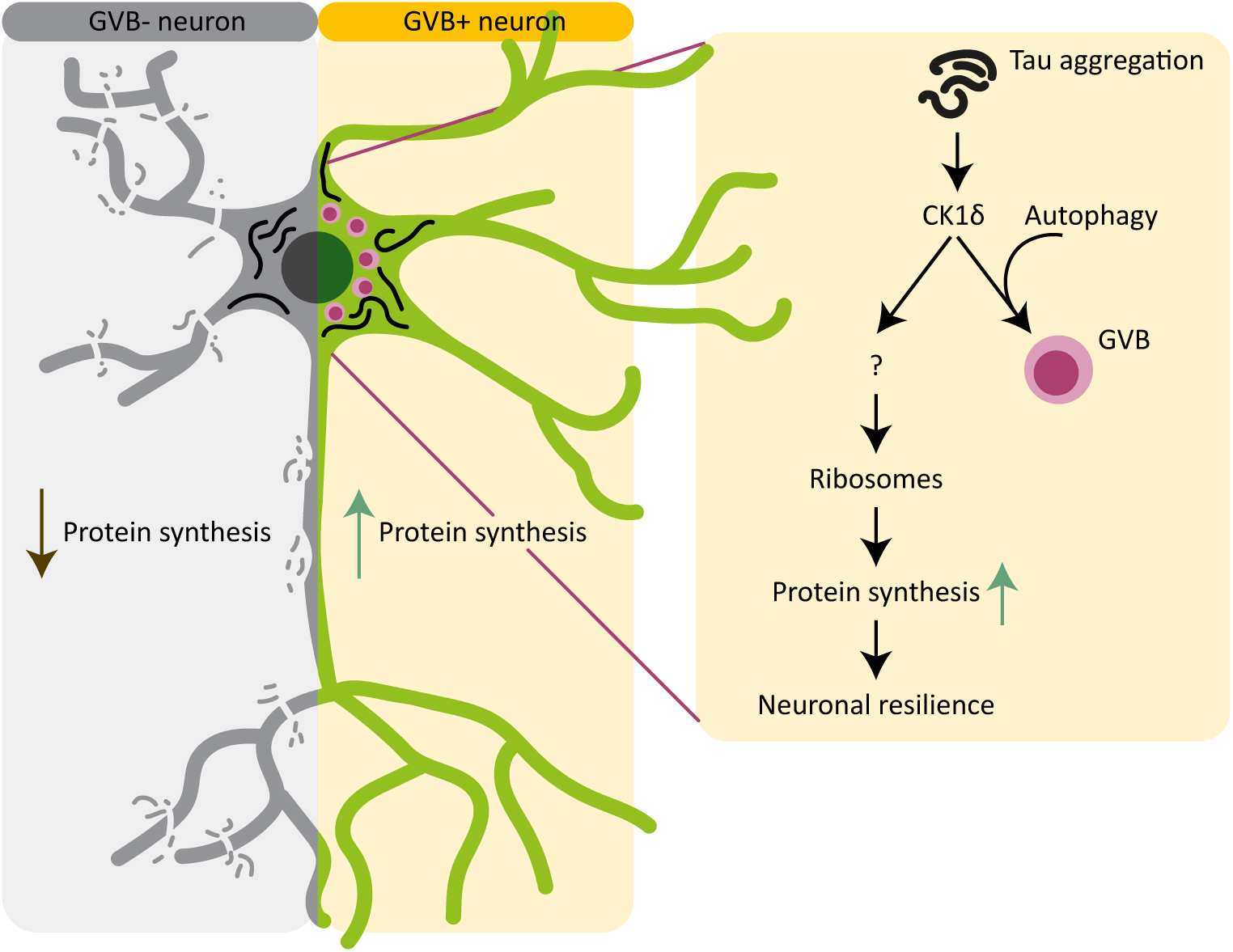

## Introduction

The progressive intraneuronal accumulation of aggregated proteins is central in the pathogenesis of neurodegenerative diseases, with tau protein in Alzheimer’s disease (AD) and frontotemporal dementias (FTD) as a prime example^1^. Neurons depend on robust protein homeostasis (proteostasis) to acquire and maintain properties required for neuronal function. Moreover, post-mitotic neurons cannot self-renew and their unique features that are acquired throughout their lifetime make them difficult to replace. Pathological tau accumulation impairs neuronal protein synthesis^2–6^, thereby strongly challenging proteostasis. Consequently, neurons must possess intrinsic mechanisms to withstand proteostatic disturbances. Granulovacuolar degeneration bodies (GVBs) are membrane-delineated structures that accumulate in the soma of a subset of neurons affected by early stages of tau pathology^7–9^, yet their functional role in tau pathogenesis remains unclear. The high number of GVB+ neurons in brains of cognitively healthy centenarians^10,11^ and the observation that the majority of surviving neurons in the AD hippocampus have GVBs^12^, suggest that the presence of GVBs may signify a protective proteostatic response. It has been hypothesized that GVBs may protect neurons by sequestering damaging proteins^9,12,13^. Alternatively, the direct connection with pathological tau accumulation could suggest that GVBs function to enhance tau degradation to restore proteostasis. However, there is no convincing evidence for the accumulation of tau in GVBs^13,14^, raising the possibility that the presence of GVBs affect neurons via an alternative mechanism. Here, we investigated the formation and functional significance of GVBs to elucidate a novel molecular mechanism underlying neuronal resilience to pathological tau-induced proteostatic disturbance.

GVBs were merely enigmatic appearances in *post-mortem* human brain tissue, until our lab recently developed experimental models of tau-induced GVB formation that allow mechanistic study^14–16^. We have demonstrated that the formation of GVBs is a neuron-specific response that is triggered by intraneuronal pathological protein assemblies^14,15^. GVBs are proteolytically active lysosomal structures that contain endo- and autosomal cargo, accumulated in one or multiple dense cores^14^. These cores are strongly immunopositive for several epitopes, including phosphorylated protein kinase R (PKR)-like endoplasmic reticulum kinase (pPERK)^13^, that are useful as markers to detect GVBs. However, it is largely unknown whether the immunopositivity of the GVB core reflects the presence of specific proteins or just protein fragments or neo-epitopes (extensively reviewed before^9,13^). To date, the serine/threonine protein kinase casein kinase 1δ (CK1δ) is the only protein validated to selectively accumulate in GVBs using direct fluorescence^14^. However, the mechanism of the formation of GVBs and the potential functional involvement of CK1δ therein is completely unknown.

In the present work, we demonstrate that neurons with GVBs (GVB+ neurons) are more resilient to pathological tau-mediated neurodegeneration. We identified CK1δ activity as a rate-limiting factor in the formation of GVBs and showed that CK1δ sequestration in the GVB requires the autophagic machinery. Neurons with GVBs have an increase in ribosomal content and are resilient to tau-induced impairment of global protein synthesis. Importantly, GVB+ neurons are resilient to the associated impairment of long-term potentiation (LTP)-regulated protein synthesis. Our results demonstrate that GVB+ neurons are protected against the proteostatic disturbance induced by tau pathology.

## Results

### GVB+ neurons are resilient to tau-induced neurodegeneration

To study the formation of GVBs in relation to tau pathology over time, we employed our extensively characterized and validated primary neuron model for seed-independent tau pathology where lentiviral transduction with FTDtau^1+2^ (2N4R human tau with the P301L/S320F FTD mutations) results in progressive pathological tau accumulation accompanied by GVB formation^15,16^ (Figure 1A). Like GVBs observed in the human brain, these GVBs contained a core positive for CK1δ- and pPERK (Figure 1B) delineated by a lysosomal membrane (Figure 1C) as previously shown^15^. 15 days after FTDtau^1+2^-transduction, around 10% of the neurons with pathological tau assemblies have formed GVBs^15,16^ (tau+/GVB+ neurons). At this point, neuronal loss or degeneration of neurites or synapses was not observed (Figures S1A-S1G), and calcium imaging analysis showed that neuronal network activity was unaffected (Figures S1H-S1L), indicating that this timepoint represents a stage in tau pathogenesis that precedes neurodegeneration. However, exposure to tau pathology for an additional week (Figure 1D) induced neuronal loss (Figure 1E). In line with studies in human brain^12^, this demonstrated that in contrast to the loss of GVB-negative (GVB-) neurons as tau pathology progresses (Figure 1F), the number of GVB+ neurons increased (Figure 1G), indicating that GVB+ neurons are more resilient to tau-induced neurodegeneration. To study the temporal dynamics of GVB formation, the percentage of GVB+ neurons was determined at different incubation periods with FTDtau^1+2^ (Figures 1H and 1I). While the accumulation of MC1- and AT8-positive tau assemblies started to plateau after 22 days of exposure to FTDtau^1+2^, the number of GVB+ neurons continued to increase up to 34% after 29 days (Figures 1I, 1J, S1M and S1N). These findings highlight that GVB formation progresses whereas GVB-neurons are lost to tau aggregation, suggesting that GVBs play a role in neuronal resilience to tau-induced neurodegeneration.

**Figure 1:**
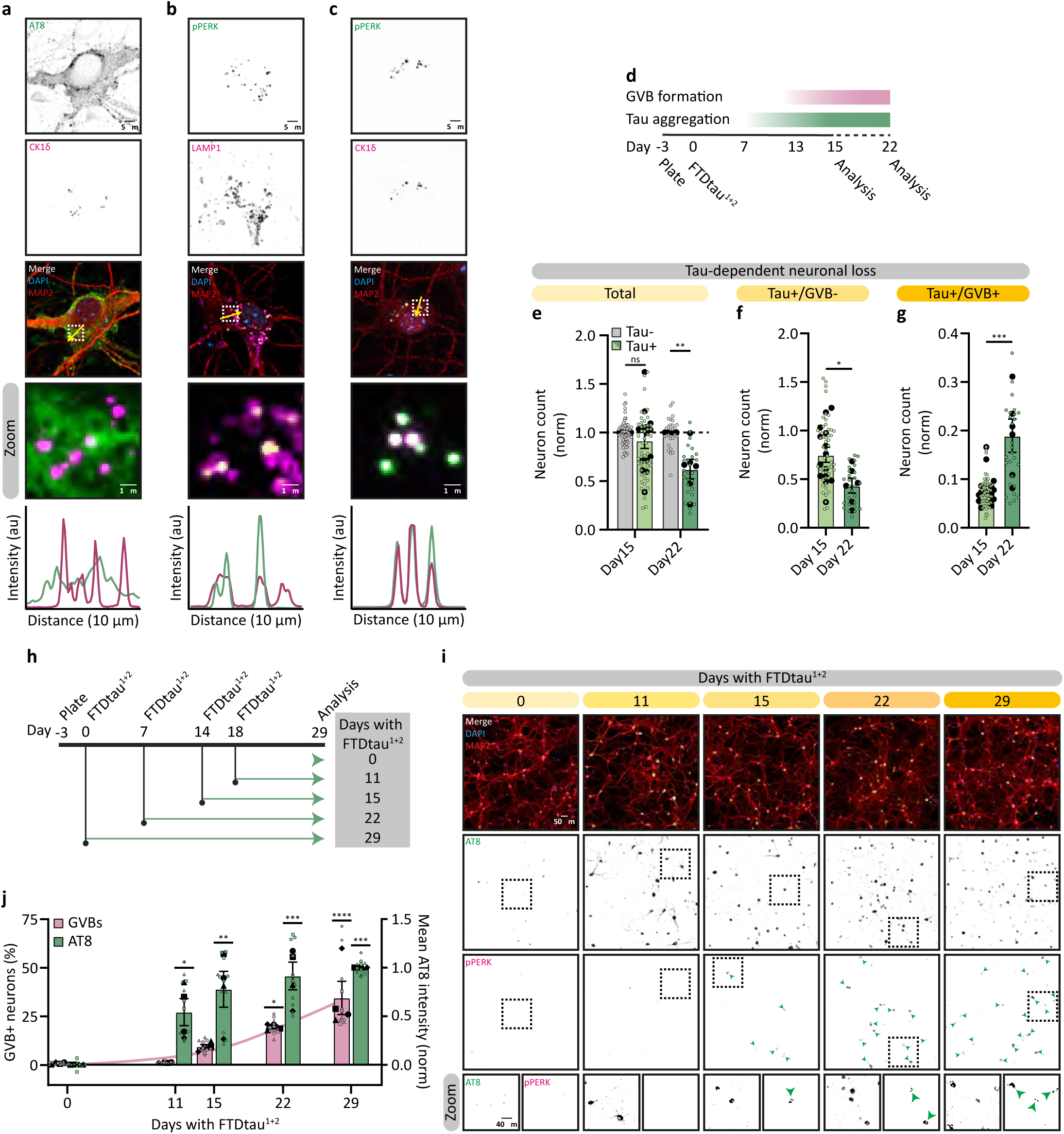
GVB+ neurons are resilient to tau-induced neurodegeneration. **A-C** Representative confocal images of tau+/GVB+ neurons analysed at day 22. Neurons were immunostained for the phosphorylated tau marker AT8 (green) and the GVB marker CK1δ (magenta) (**A**), the GVB markers pPERK (green) and CK1δ (magenta) (**B**), pPERK (green) and lysosomal membrane marker LAMP1 (magenta) (**C**) and the dendrite marker MAP2 (red). The zoomed regions are indicated by dashed white squares and the yellow arrow represents the location of the line intensity profiles. **D** Schematic of the experimental timeline indicating the time of transduction with FTDtau^1+2^, start of tau aggregation and GVB formation. Analysis was performed at day 15 or 22. **E-G** Quantification of tau-dependent neuron number (tau+; **E**) in tau+/GVB- (**F**) or tau+/GVB+ (**G**) neurons analysed at day 15 or day 22, normalised to the mean number of tau-neurons per timepoint. For day 15, N=14 and n=54 (tau-) and 52 (tau+/GVB- and tau+/GVB+). For day 22, N=6 and n=27 (all). For visual clarity, the biological replicates are not distinguished. A nested t-test was used. **H** Schematic of the protocol used to assess the progression of tau aggregation and GVB+ formation by transduction with FTDtau^1+2^ at the indicated points. **I** Representative high-content microscopy images at different FTDtau^1+2^ exposure durations as indicated. Neurons were immunostained for AT8 (green), MAP2 (red) and pPERK (magenta). The zoomed regions are indicated by dashed black squares. Green arrowheads indicate GVB+ neurons. **J** Quantification of the percentage GVB+ neurons and the mean AT8 immunofluorescence intensity normalised to 29 days FTDtau^1+2^ exposure (1) and untransduced (0). N=4 and n=10, 10, 12, 12 and 11 for 0, 11, 15, 22 and 29 days with FTDtau^1+2^, respectively. A nested one-way ANOVA followed by a Dunnett’s post-hoc test was used. **K** Representative confocal images of tau+/GVB+ neurons treated with lysosomal inhibitors E64D and PepstatinA (PepA) for 24h analysed at day 15. Neurons were immunostained for AT8 (green), LAMP1 (red), pPERK (magenta) and MAP2 (blue). **l** Representative confocal images of tau+/GVB+ neurons analysed at day 15. Neurons were treated with Proteostat (red) to detect aggregated proteins and immunostained for AT8 (green), pPERK (magenta) and MAP2 (blue). Nuclei were visualised by DAPI (blue) and separate channels are shown in greyscale (**A-C, I, K, L**). Data are presented as mean ± SEM. *p*-values are indicated: \**p* < 0.05, \*\**p* < 0.01, \*\*\**p* < 0.001, \*\*\*\**p* < 0.0001 and ns: not significant.

### CK1δ activity is a key regulator of GVB formation

To investigate the molecular mechanism associated with the formation of GVBs, the kinase CK1δ is a prime candidate because it selectively accumulates in GVBs^14^. To study the role of CK1δ, tau+ neurons were treated with the CK1δ/ε-specific ATP-competitive kinase inhibitor PF670462^17,18^ (CK1δ_i_). We studied the effect of CK1δ_i_ 15 days after FTDtau^1+2^-transduction. CK1δ_i_ treatment was initiated either before or after the first GVBs are detected at day 13^15^ (Figures 2A and 2B). CK1δ_i_ did not induce toxicity or negatively affect neuronal morphology or synapse number (Figures S2A-S2M). Using pPERK as a GVB marker, we found that CK1δ_i_ treatment for 48h or 8d reduced the number of GVB+ neurons to 33% and 4% of control, respectively (Figures 2A and 2C). This shows that the formation of GVBs is reduced by CK1δ_i_ when initiated before or during the first GVBs occur. In contrast, CK1δ_i_ treatment for 24h or less did not significantly change the number of GVB+ neurons (Figures 2A and 2C). To test if this was due to the shorter incubation period or if CK1δ_i_ has no effect once GVB formation is in progress, tau pathology exposure was extended to 22 days, with treatment starting well after the onset of GVB formation (Figure S2N). The number of GVB+ neurons was also strongly reduced upon CK1δ_i_ treatment after 48h (63% of control) and 8d (12% of control) (Figure S2O), indicating that CK1δ is required both for the initiation and progression of GVB formation. To exclude that the effect of CK1δ_i_ was due to altered phosphorylation of GVB-cargo that may affect the detection of phospho-epitopes like pPERK, we tested the effect of CK1δ_i_ using a non-phospho GVB marker that has been identified in AD post-mortem brain, GOLGINA4^19,20^. GOLGINA4 was validated as a bona fide GVB marker in our experimental model as well (Figures S3A and S3B). CK1δ_i_ treatment strongly reduced GOLGINA4-detected GVB+ neurons to 27% of control (Figure S3C), confirming that CK1δ_i_ decreases GVB formation. CK1δ is reported to phosphorylate tau *in vitro*^21,22^, therefore we assessed pathological tau accumulation as previously described^15,16,23^. Treatment with CK1δ_i_ for 48h or 8d did not affect the level of MC1-positive FTDtau^1+2^ accumulation (Figures S3D and S3F) and the phospho-epitope-independent direct fluorescence intensity of MeOH-insoluble FTDtau^1+2^-GFP (Figures S3E and S3G). This indicates that the effect of inhibition of CK1δ activity on the number of GVB+ neurons is not caused by an indirect effect on tau pathology. Together, these data collectively place CK1δ upstream in the pathway driving GVB formation.

**Figure 2:**
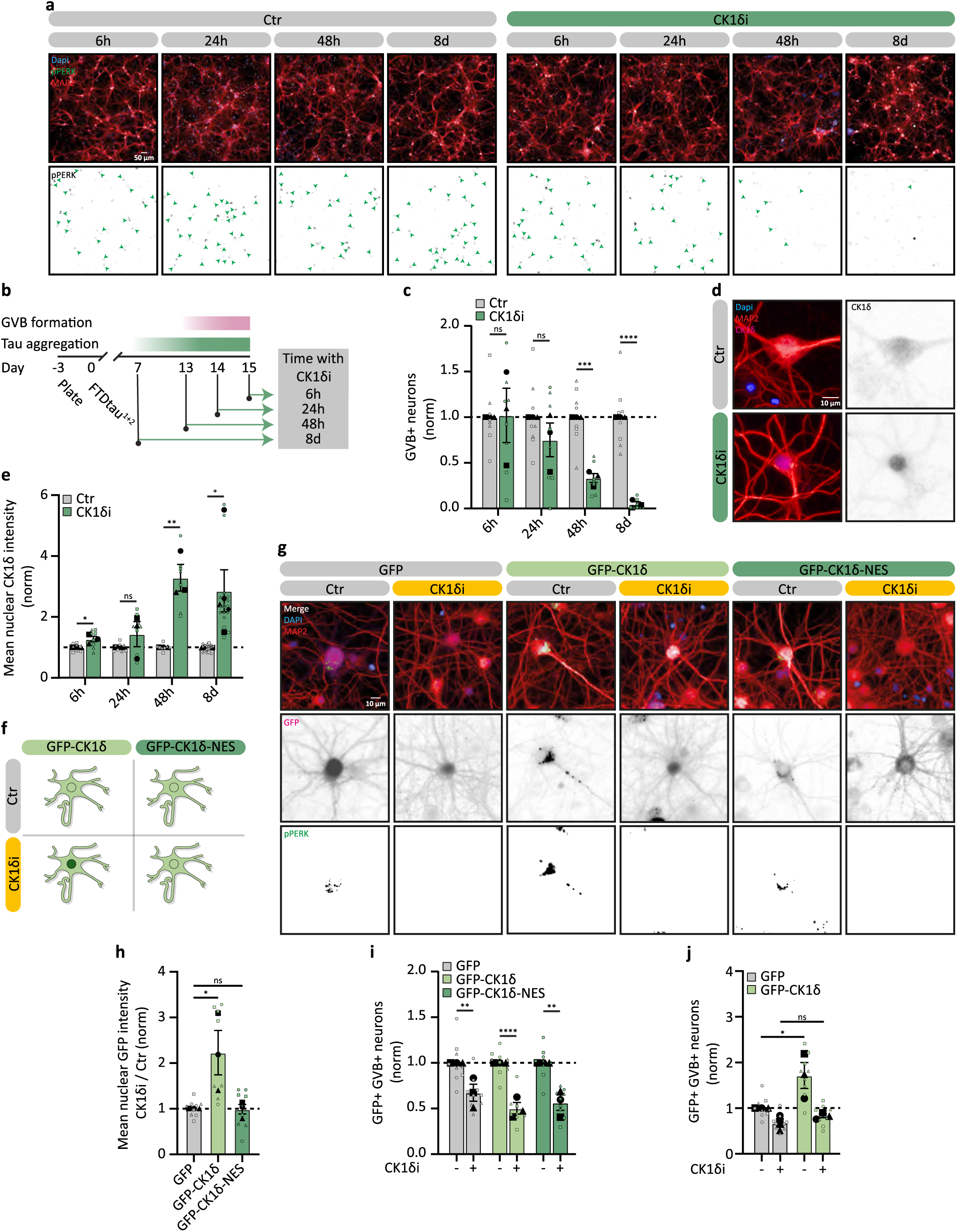
CK1δ is a key regulator for the formation of GVBs. **A** Representative high-content microscopy images of tau+ neurons without or with CK1δ_i_ for 6h, 24h, 48h or 8d analysed at day 15. Neurons were immunostained for the dendrite marker MAP2 (red) and the GVB marker pPERK (green). Green arrowheads indicate single GVB+ neurons. **B** Schematic of the timeline of CK1δ_i_. **C** Quantification of the percentage of pPERK+ GVB+ neurons upon CK1δ_i_ treatment analysed at day 15 normalised to untreated per timepoint. N=3 and n=8 and 8 (6h, 24h, 8d), 8 and 7 (48h), for Ctr and CK1δ_i_, respectively. A nested t-test was used. **D** Representative high-content microscopy images of tau+ neurons treated without or with CK1δ_i_ for 48h analysed at day 15. Neurons were immunostained for MAP2 (red) and CK1δ (magenta). **E** Quantification of mean nuclear CK1δ intensity upon CK1δ_i_ treatment analysed at day 15 normalised to untreated per timepoint. N=3 and n=8 and 8 (6h, 24h), 8 and 7 (48h) and N=5 and n=17 and 17 (8d), for Ctr and CK1δ_i_, respectively. A nested t-test was used. **F** Schematic representation of subcellular localisation of GFP-CK1δ and GFP-CK1δ-NES in the absence and presence of CK1δ_i_. **G** Representative high-content microscopy images of tau+ neurons transduced with GFP, GFP-CK1δ or GFP-CK1δ-NES without or with CK1δ_i_ for 48h analysed at day 22. Shown are GFP signal (magenta) and immunofluorescent signal for MAP2 (red) and pPERK (green). **H** Quantification of mean nuclear GFP signal of neurons transduced with GFP, GFP-CK1δ or GFP-CK1δ-NES upon 48h of CK1δ_i_ treatment in GFP-selected neurons analysed at day 22. First, the nuclear GFP intensity was normalised per group upon CK1δ_i_, after which it was normalised to the response in GFP-transduced neurons. N=3 and n=9 (GFP), 8 (GFP-CK1δ), 9 (GFP-CK1δ-NES). A nested one-way ANOVA followed by a Dunnett’s post-hoc test was used. **I** Quantification of the percentage of pPERK+ GVB+ neurons of neurons transduced with GFP, GFP-CK1δ or GFP-CK1δ-NES upon 48h of CK1δ_i_ treatment in GFP-selected neurons analysed at day 22 normalised to untreated per group. N=3 and n=9 and 9 (GFP), 9 and 8 (GFP-CK1δ), 9 and 9 (GFP-CK1δ-NES), for Ctr and CK1δ_i_, respectively. A nested t-test was used. **J** Quantification of the percentage of pPERK+ GVB+ neurons transduced with GFP or GFP-CK1δ, upon 48h of CK1δ_i_ treatment in GFP-selected neurons analysed at day 22 and normalised to untreated GFP-transduced neurons. N=3 and n=9 and 9 (GFP), 9 and 8 (GFP-CK1δ), for Ctr and CK1δ_i,_ respectively. A nested one-way ANOVA followed by a Sidak’s post-hoc test was used. Nuclei were visualised by DAPI (blue) and separate channels are shown in greyscale (**A, D, G**). Data are presented as mean ± SEM. *p*-values are indicated: \**p* < 0.05, \*\**p* < 0.01, \*\*\**p* < 0.001, \*\*\*\**p* < 0.0001 and ns: not significant.

To further confirm the key role of CK1δ in GVB generation, shRNA-mediated KD was employed to reduce CK1δ protein levels (Figures S4A-S4C). *shCsnk1d* significantly reduced the percentage of GVB+ neurons to 52% of control (Figures S4D and S4E). Because of the high toxicity associated with *shCsnk1d*-treatment (Figure S4F), overexpression of a *shRNA*-resistant CK1δ variant (GFP-CK1δ_RES_) was used to distinguish the effects of CK1δ KD on the formation of GVBs from its toxic effects (Figures S4G-S4J). This was sufficient to rescue the toxic effect of CK1δ KD (Figure S4K). Moreover, overexpression of GFP-CK1δ_RES_ rescued the inhibition of GVB accumulation induced by *shCsnk1d* (Figure S4L), further supporting an essential role for CK1δ in the formation of GVBs.

Next, we investigated if regulation of CK1δ levels or activity underlies its role in GVB formation. Tau aggregation did not alter the total protein level of CK1δ (Figures S5A and S5B) with no difference in somatic CK1δ intensity between GVB-and GVB+ neurons (Figure S5C). In contrast, treatment with CK1δ_i_ strongly increased CK1δ levels, independent of the presence of tau (Figures S5A and S5B). Moreover, CK1δ_i_ increased nuclear CK1δ levels (Figures 2D, 2E and S2P), in line with a previous study that showed localisation of inactive CK1δ to the nucleus^24^. To exclude that the effect of CK1δ_i_ on GVBs was indirectly caused by a changed balance in cytosolic-to-nuclear CK1δ levels, we employed GFP-CK1δ with a nuclear export signal (GFP-CK1δ-NES) to force its cytosolic localisation (Figure 2F). GFP-CK1δ-NES selectively localised to GVBs (Figures S5D and S5E), indicating that the NES sequence does not interfere with the localisation of CK1δ to GVBs. Upon treatment with CK1δ_i_, GFP-CK1δ relocalised to the nucleus like endogenous CK1δ, whereas -as expected-the localisation of GFP-CK1δ-NES was not changed (Figures 2G and 2H). However, GFP-CK1δ-NES was not able to rescue the CK1δ_i_-induced decrease in number of GVB+ neurons (Figures 2I), indicating that the activity of CK1δ directly affects the formation of GVBs. To further support the direct involvement of CK1δ activity in GVB formation, GFP or GFP-CK1δ^14^ was overexpressed in tau+ neurons (Figure 2G). GFP-CK1δ+ neurons were 1.72 times more likely to form GVBs than GFP+ neurons (Figure 2J). Moreover, the effect of GFP-CK1δ overexpression on GVBs was lost when neurons were treated with CK1δ_i_ (Figure 2J). This demonstrated that the activity of CK1δ is required to generate GVB+ neurons.

### CK1δ accumulation in GVBs marks the appearance of GVB+ neurons

To better understand the connection between the role of CK1δ activity in GVB formation and its accumulation inside GVBs, we first investigated whether the effect of CK1δ_i_ on GVB+ neurons is binary (GVB+ or GVB-neuron) or more continuous (affecting the number of GVBs per neuron). Using high-resolution confocal microscopy, reduced GVB content per neuron upon CK1δ_i_ treatment was observed (Figure 3A). Quantification by high-throughput microscopy confirmed that inhibition of CK1δ decreased the total area of GVBs within a single neuron, but not the size or the GVB over cytosol (G/C) ratio of CK1δ fluorescence intensity of individual GVBs (Figures 3B-3D). These data indicate that CK1δ activity stimulates the formation of GVBs also after the first GVBs have formed. To investigate the role of CK1δ activity in CK1δ localisation to GVBs more directly, a kinase-dead K38M CK1δ mutant was employed (Figure S5E)^24,25^. GFP-CK1δ-K38M selectively accumulates in GVBs (Figure 3E). This indicates that although CK1δ activity is necessary for GVB formation, the activity of an individual CK1δ molecule (*in cis* activity) is not required for its specific accumulation in the core of GVBs.

**Figure 3:**
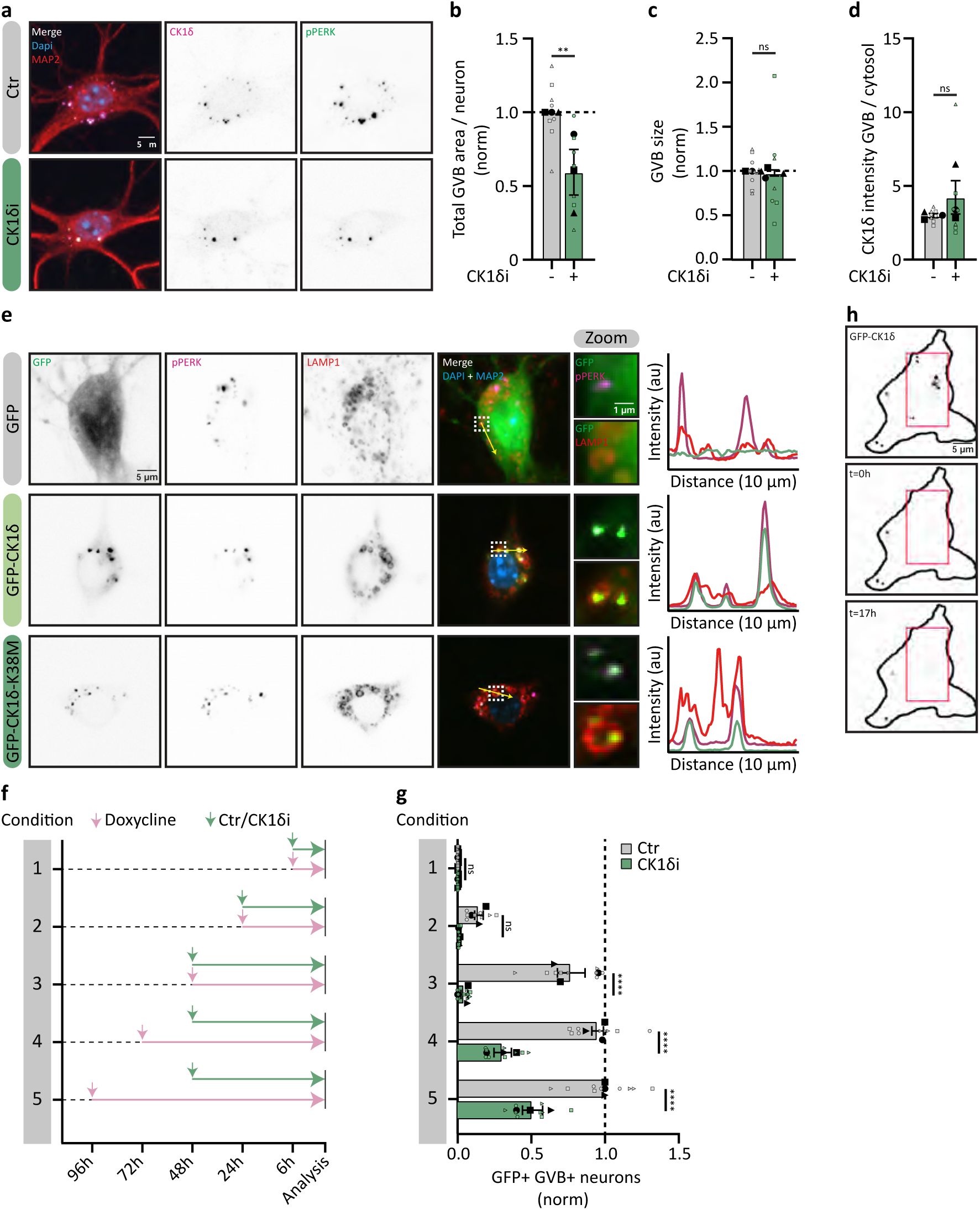
CK1δ accumulation in GVBs marks the appearance of GVBs. **A** Representative confocal images of tau+/GVB+ neurons without or with CK1δ_i_ for 48h analysed at day 22 (maximum intensity projected Z-stacks). Neurons were immunostained for the GVB markers CK1δ (magenta) and pPERK (green) and the dendrite marker MAP2 (red). **B-D** High-throughput quantification of pPERK+ GVB-area per neuron (**B**), single GVB size (**C**) and the ratio of CK1δ intensity in the GVB over the cytosol (**D**) of tau+/GVB+ neurons without or with CK1δ_i_ for 48h analysed at day 22. Data were normalised to control (**B, C**). N=3 and n=8 and 7, for Ctr and CK1δ_i_, respectively. A nested t-test was used. **E** Representative confocal images of tau+/GVB+ neurons analysed at day 22. Neurons were transduced with GFP, GFP-CK1δ or GFP-CK1δ-K38M. The neurons were immunostained for pPERK (magenta), the lysosomal membrane marker LAMP1 (red) and MAP2 (blue). The zoomed regions are indicated by dashed white squares and the yellow arrow represents the location of the line intensity profiles. **F** Schematic overview of tau+ neurons transduced with Dox-GFP-CK1δ treated with doxycycline (Dox) for 6h, 24h, 48h, 72h or 96h as indicated by the pink arrow and without or with CK1δ_i_ for 6h, 24h or 48h indicated by the green arrow. **G** Quantification of the percentage of GFP+/GVB+ neurons treated with Dox and without or with CK1δ_i_ according to the conditions specified in **F** normalised to control neurons treated for 96h with Dox analysed at day 22. GVBs were defined using Dox-GFP-CK1δ as the GVB-marker. N=3 and n=8 and 8 (condition 2), 9 and 9 (the other conditions), Ctr and CK1δ_i_, respectively. A nested one-way ANOVA followed by a Sidak’s post-hoc test was used. **H** Representative live confocal images of tau+/GVB+ neurons transduced with GFP-CK1δ and followed over time after FRAP. Shown are cells directly after FRAP (t=0h) and at the end of the experiment (t=17h) is shown. The red square indicates the region that was exposed to FRAP. Nuclei were visualised by DAPI (blue) and separate channels are shown in greyscale (**A, E**). Data are presented as mean ± SEM. *p*-values are indicated: \*\**p* < 0.01, \*\*\*\**p* < 0.0001 and ns: not significant.

To study the dynamics of CK1δ accumulation in GVBs, we used a doxycyclin (Dox)-inducible expression system for GFP-CK1δ (Dox-GFP-CK1δ; Figures S6A and S6B). Dox-GFP-CK1δ was first detected in GVBs 24h after induction and strongly increased at 48h, after which it started to plateau (Figures 3F and 3G), indicating that a steady-state of newly labelled GVBs is obtained at this point. In the presence of CK1δ_i_ (condition 1-3, Figure 3F), the formation of neurons with Dox-GFP-CK1δ+ GVBs was completely prevented (Figure 3G). Because CK1δ activity *in cis* is not required for GVB localisation (Figure 3E), this suggests that the targeting of CK1δ to the core of GVBs is directly coupled to the formation of GVBs. To formally exclude the possibility that rather than inhibiting the formation of new GVBs, CK1δ_i_ stimulates the turnover of already formed GVBs, Dox-GFP-CK1δ expression was induced 24h or 48h before CK1δ_i_ treatment (condition 4-5, Figure 3F). CK1δ_i_ treatment reduced the number of Dox-GFP-CK1δ+/GVB+ neurons, but did not completely block it (Figure 3G). This indicates that the remaining Dox-GFP-CK1δ+/GVB+ neurons were formed before CK1δ_i_ was added. Consequently, we were able to calculate the stability of GVBs. Since CK1δi blocks new GVB formation, the observed GVB+ neurons (Figure S2O) existed before treatment. The difference between expected (Figure 1H) and observed GVB+ neurons (Figure S2O) was used to calculate an average t₁_/_₂ of 5.4 days (Figure S6C). This is supported by live cell microscopy of GFP-CK1δ-labeled GVBs, demonstrating that GVBs were relatively stable for at least 9h (Video S1), indicating a low-turnover rate. Furthermore, using fluorescence recovery after photobleaching (FRAP) imaging, we observed that no new CK1δ is incorporated into the pre-existing GVBs over a span of ±17h (Figure 3H). These data collectively identify CK1δ as an upstream regulator of the pathway that makes neurons GVB-positive and indicate that CK1δ is targeted to the GVB core during this process. Given the stability of the CK1δ accumulations in GVBs, they reflect the initiation of the GVB condition in the neuron.

### Cargo accumulaton in GVBs requires autophagy

The most likely way for proteins like CK1δ to end up in the GVB core is via autophagy. To explore the role of autophagy in GVB formation, tau+ neurons were treated with Sar405 to block VPS34 (Figures 4A and S7A). This efficiently inhibited autophagic flux, as indicated by strongly increased levels of P62 (Figure S7B). In addition, the number of GVB+ neurons gradually decreased over time down to 22% of control after 8d treatment (Figures 4A and 4B). To further support the involvement of autophagy in GVB formation, knock-down (KD) of *Fip200* and *Atg5*, that act after autophagy initiation, was performed (Figure S7A). *Fip200* as well as *Atg5* shRNAs^26,27^ efficiently reduced the protein levels of their targets (Figures S7C, S7D, S7F and S7G) and both increased P62 levels (Figures S7E and S7H), showing inhibition of autophagy. All shRNAs reduced the number of GVB+ neurons (*shFip200* to 39% and *shAtg5* to 25% of control) (Figures 4C-4F). To test the effect of activation of autophagy, we employed the mTOR inhibitors Torin1 and rapamycin for 48h (Figure S7A). Both treatments strongly reduced 4E-BP1 phosphorylation, showing efficient inhibition of mTOR activity (Figures S7I and SK). In contrast, the number of GVB+ neurons was not affected by either treatment (Figures S7J and S7L). Together, these results demonstrate that GVB formation is dependent on the autophagic machinery and flux but is not stimulated by increasing autophagic activity.

**Figure 4:**
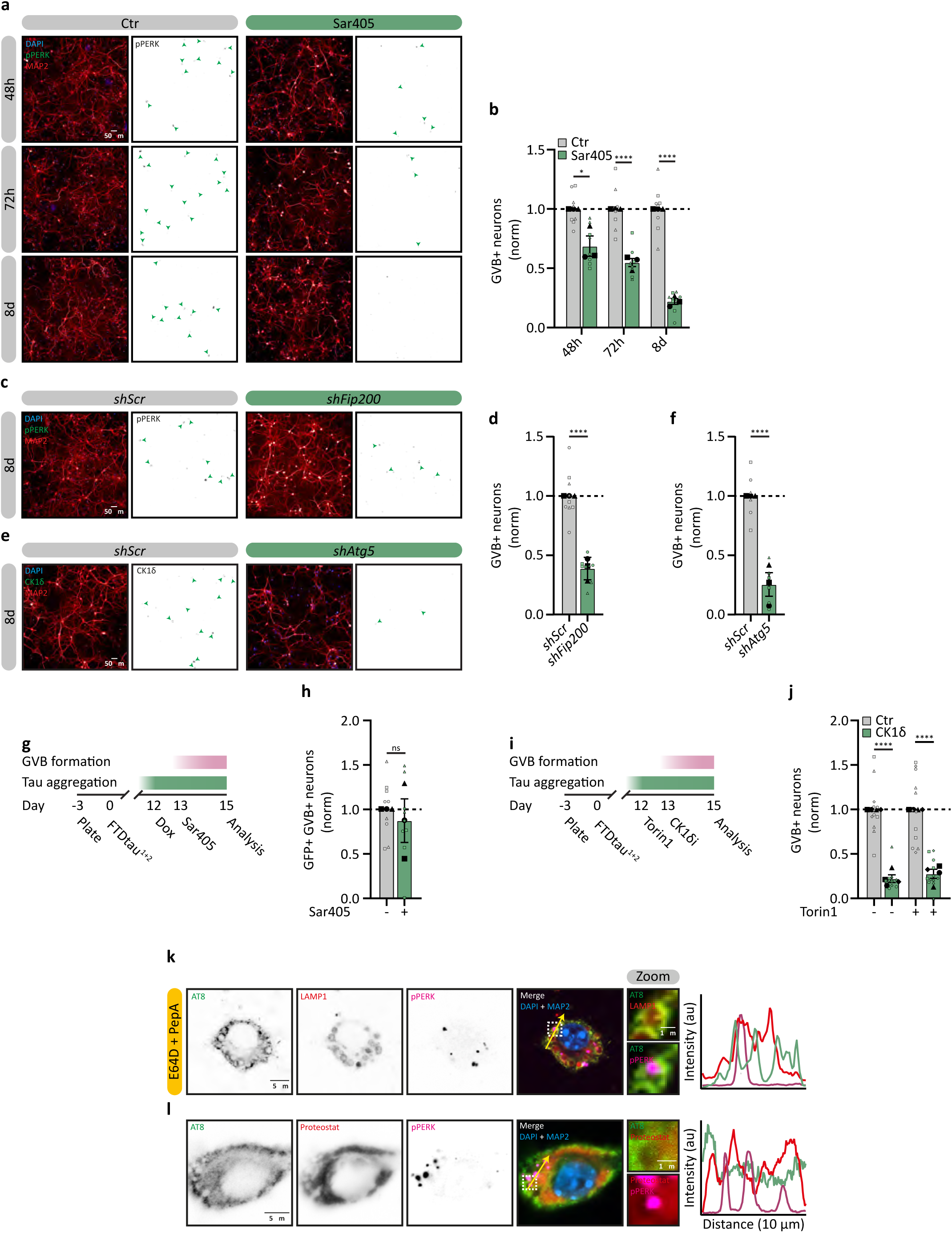
CK1δ sequestration into the GVB requires basal autophagy. **A** Representative high-content microscopy images of tau+ neurons treated without or with the VPS34 chemical inhibitor Sar405 for 48h, 72h or 8d analysed at day 15. Neurons were immunostained for the GVB marker pPERK (green) and the dendrite marker MAP2 (red). **B** Quantification of the percentage of pPERK+ GVB+ neurons upon Sar405 treatment normalised to untreated per timepoint at day 15. N=3 and n=9 and 8 (48h), n=8 and 8 (72h), n=8 and 9 (8d), Ctr and Sar405, respectively. A nested t-test was used. **C, E** Representative high-content microscopy images of tau+ neurons transduced with a control (*shScr*), shRNA targeting *Fip200* (*shFip200*; **C**) or *Atg5* (*shAtg5;* **E**) 8d before analysis at day 15. Neurons were immunostained for MAP2 (red) and pPERK (**C**) or CK1δ (**E**) (green). **D, F** Quantification of the the percentage of pPERK+ (**D**) or CK1δ+ (**F**) GVB+ neurons upon *Fip200* (**D**) and *Atg5* knock-down (KD) (**F**) normalised to *shScr*. N=3 and n=9 and 8 (**D**), n=6 and 6 (**F**), *shScr* and *shFip200* or *shAtg5*, respectively. A nested t-test was used. **G** Schematic of the timeline of induction of CK1δ expression (Dox) and inhibition of autophagy (Sar405). **H** Quantification of the percentage of GFP+/GVB+ neurons transduced with Dox-GFP-CK1δ and treated with Dox for 72h, upon 48h of Sar405 treatment, analysed at day 15 and normalised to untreated. GVBs were defined using Dox-GFP-CK1δ as the GVB-marker. N=3 and n=9. A nested t-test was used. **I** Schematic of the timeline of induction of autophagy (Torin1) and CK1δi. **J** Quantification of the percentage of pPERK+ GVB+ neurons upon CK1δi treatment without or with prior Torin1 treatment analysed at day 15 normalised to untreated. N=4 and n=11 (CK1δ_i_ only) or 12 (the rest). A nested t-test was used. Nuclei were visualised by DAPI (blue) and separate channels are shown in greyscale (**A, C, E**). Data are presented as mean ± SEM. *p*-values are indicated: \**p* < 0.05, \*\*\*\**p* < 0.0001 and ns: not significant.

These data support a role for autophagy in the sequestration of proteins, like CK1δ, in the GVB core after the initiation of the pathway. To investigate whether autophagy is also an upstream regulator of the pathway that makes neurons GVB+, we studied the connection between CK1δ and autophagy in GVB formation. To this end, we used the Dox-inducible expression system for GFP-CK1δ. Inhibition of autophagy with Sar405 after increasing CK1δ levels by the addition of Dox did not affect CK1δ accumulation in GVBs (Figures 4G and 4H), In line with this, stimulation of autophagy with Torin1 before treatment with CK1δ_i_ (Figure 4I), did not rescue the CK1δ_i_-induced reduction in GVB+ neurons (Figure 4J). The combined results place CK1δ activity upstream in the pathway leading to GVB+ neurons and suggest that autophagy sequesters CK1δ in the GVB after it has performed its role in the process.

Pathological tau does not accumulate in the GVB in the human brain and experimental models^9,13^. To exclude that GVBs rapidly degrade tau, thereby precluding the detection of tau inside the GVB, lysosomal degradation was inhibited using E64D and Pepstatin A. This strongly increased in the accumulation of P62 (Figure S8A), confirming effective lysosomal inhibition. However, this treatment did not result in the accumulation of AT8-positive tau in the GVB (Figure 4K). In addition, whereas intraneuronal tau pathology was positive for the aggregate-binding dye Proteostat, no positive signal was observed in GVBs (Figure 4L). This suggests that GVBs do not play a key role in pathological tau degradation.

### GVB+ neurons are resilient to tau-induced protein synthesis impairment

To investigate alternative potential functional implications of the presence of GVBs, early proteome changes induced by tau pathology at day 15 were determined (Figure 5A and Table S1). Gene ontology (GO) analysis indicated that the upregulated functional processes primarily involved pathways affecting the synapse (Figure 5B) in line with the tau/GVB model representing an early stage preceding neurodegeneration (Figures S1A-S1L). The most significantly downregulated functional protein groups are solute carrier membrane transport proteins, predominantly amino acid transporters (Figure 5B). This may impact the availability of amino acids for the biosynthesis of proteins and is in line with reduced neuronal protein synthesis upon pathological tau accumulation^2–6^. To directly determine *de novo* protein synthesis in relation to tau pathology and GVB state, we employed the SUnSET puromycin labelling assay^28,29^ combined with quantitative high-content automated microscopy to analyse tau-and tau+ neurons at day 22. We developed a script to differentiate between tau+/GVB-and tau+/GVB+ neurons within the tau+ population. Tau+/GVB-neurons, which constitute the majority of tau+ neurons, displayed a pronounced reduction in protein synthesis rates of 31% compared to tau-neurons (Figures 5C and 5D). In contrast, tau+/GVB+ neurons retained protein synthesis rates similar to those of tau-neurons (Figures 5C and 5D).

**Figure 5:**
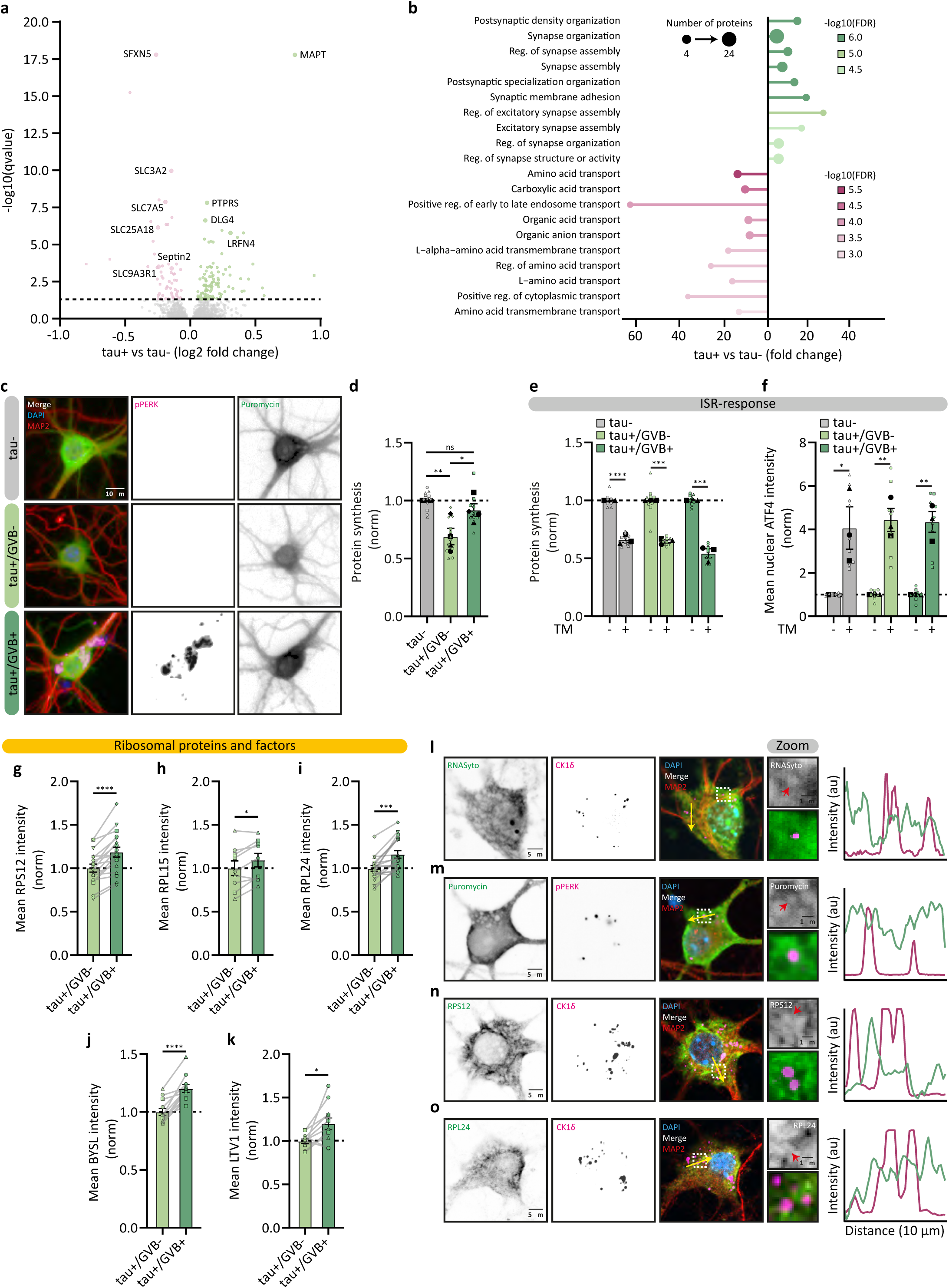
GVB+ neurons are resilient to tau-induced protein synthesis impairment. **A** Volcano plot showing significantly upregulated (green) and downregulated (pink) proteins in tau+ neurons (q-value < 0.05) analysed at day 15. See Methods for details. See Table S1. **B** Most significant terms resulting from the GO enrichment analysis of significantly upregulated (green) and downregulated (pink) proteins in tau+ neurons. The circle size at the tip of the bar represents the number of significant proteins involved in the process and the colour intensity is indicative of the significance of the GO term. See Methods for details. See Table S1. **C** Representative high-content microscopy images showing tau-, tau+/GVB- and tau+/GVB+ neurons analysed at day 22. Neurons were treated for 15min with puromycin and were immunostained for puromycinilated proteins (green), the dendrite marker MAP2 (red) and the GVB marker pPERK (magenta). **D** Quantification of puromycin intensity as a measure for *de novo* protein synthesis in tau-, tau+/GVB- and tau+/GVB+ (pPERK based) neurons analysed at day 22, normalised to tau-neurons. N=4 and n=12. A nested one-way ANOVA followed by a Tukey’s post-hoc test was used. **E** Quantification of puromycin intensity as a measure for *de novo* protein synthesis in tau-, tau+/GVB- and tau+/GVB+ (pPERK based) neurons treated with TM for 24h analysed at day 22, normalised to each group. N=3 and n= 8 and 9 (tau-), 9 and 9 (tau+/GVB-), 9 and 9 (tau+/GVB+), for Ctr and TM, respectively. A nested t-test was used. **F** Quantification of mean nuclear ATF4 intensity in tau-, tau+/GVB- and tau+/GVB+ (CK1δ based) neurons treated with TM for 24h analysed at day 22, normalised to each group. N=3 and n=9. A nested t-test was used. **G-K** Quantification of somatic intensities in tau+/GVB- and tau+/GVB+ (CK1δ based) neurons analysed at day 22 for 40S ribosomal protein S12 (RPS12; **G**), 60S ribosomal protein L15 (RPL15; **H**), 60S ribosomal protein L24 (RPL24**; I**) and ribosome biogenesis factors BYSL (**J**) and LTV1 (**K**). RPS12 (**G**): N=6 and n=18. RPL15 (**H**): N=3 and n=9. RPL24 (**I**): N=6 and n=18. BYSL (**J**): N=4 and n=11. LTV1 (**K**): N=4 and n=11. A paired t-test was used. **L-O**, Representative confocal images of tau+/GVB+ neurons analysed at day 22. Neurons were treated with SYTO RNASelect to visualise RNA for 4h (green) and immunostained for CK1δ (magenta) (**L**), puromycin (green) and pPERK (magenta) (**M**), RPS12 (green) and CK1δ (magenta) (**N**) and RPL24 (green) and CK1δ (magenta) (**O**) and MAP2 (red). Nuclei were visualised by DAPI (blue). The zoomed regions are indicated by dashed white squares and the yellow arrow represents the location of the line intensity profiles. The red arrow points to a single GVB core. Nuclei were visualised by DAPI (blue) and separate channels are shown in greyscales (**C, L-O**). Data are presented as mean ± SEM. *p*-values are indicated: \**p* < 0.05, \*\**p* < 0.01, \*\*\**p* < 0.001, \*\*\*\**p* < 0.0001 and ns: not significant.

Next, we investigated whether these differences in protein synthesis were mediated by transient adaptations to proteostatic stress, of which the integrated stress response (ISR) is a key regulatory pathway. To test if the GVB+ neurons actively maintained protein synthesis rates or were less responsive to ISR-induced protein synthesis reduction, neurons were treated with the ISR inducer tunicamycin (TM). TM treatment resulted in a strong reduction in protein synthesis and increased nuclear ATF4 intensity independent of the presence of tau pathology or GVBs (Figures 5E and 5F). This shows that tau-, GVB- and GVB+ neurons can all dynamically adjust their protein synthesis levels in response to stress. Treatment with the ISR inhibitor ISRIB did not affect protein synthesis rates in GVB- or GVB+ neurons (Figure S9B), further supporting that the ISR is not involved in GVB-associated protein synthesis resilience. Protein synthesis can also be transiently reduced by the mTOR pathway. However, there was no difference in the phosphorylation of the mTOR substrate 4E-BP1 neurons without and with tau pathology, suggesting that the mTOR pathway is not activated in the pool of tau+ neurons (Figures S9C and S9D). Also, treatment with pharmacological inhibitors of the mTOR- and Angiogenin-mediated stress pathways did not affect protein synthesis rates in tau+/GVB- or tau+/GVB+ neurons (Figures S9E and S9F). These data collectively show that transient stress pathways are not directly involved in the tau-induced protein synthesis reduction or the resilience against it in GVB+ neurons.

To determine whether adaptations in the protein synthesis machinery itself underlie the resilience, we analysed the levels of ribosomal proteins and biogenesis factors in tau+ neurons. The levels of ribosomal proteins RPL24, RPL15 and RPS12 were increased in tau+/GVB+ neurons compared to tau+/GVB-neurons by 16%, 9% and 18%, respectively (Figures 5H-5J). Also, the levels of ribosomal assembly factors BYSL and LTV were 20% higher in tau+/GVB+ neurons (Figures 5K and 5L). To investigate whether the GVB itself functions as a localised hub for protein synthesis, we assessed the colocalisation of GVBs with key components of the translation machinery using confocal microscopy. We stained for RNA, using RNASyto Green, nascent proteins via puromycin pulse labelling and RPS12 and RPL24 (Figures 5M-5P). No overlap was observed between these markers and GVBs, making it unlikely that GVBs themselves directly contribute to protein synthesis. These results suggest that GVB+ neurons elicit a response to ameliorate reduced protein synthesis upon tau aggregation that includes increased ribosomal content.

### GVB+ neurons retain the capacity to control LTP-regulated protein synthesis in the presence of tau pathology

We showed that GVB+ neurons are resilient to tau-induced neurodegeneration (Figures 1E-1G). To investigate whether the increased survival of GVB+ neurons is related to protein synthesis capacity, protein synthesis was inhibited using 48h of anisomycin treatment (Figure 6A), which reduced protein synthesis to a similar extent in GVB- and GVB+ neurons, 65% and 66% respectively (Figure 6B). However, while anisomycin treatment resulted in synthetic lethality with tau pathology in GVB-neurons (28% neuron loss), GVB+ neuron numbers were unaffected (Figure 6C). These results suggest that GVB+ neurons are more resilient to toxicity induced by pharmacological protein synthesis inhibition.

**Figure 6:**
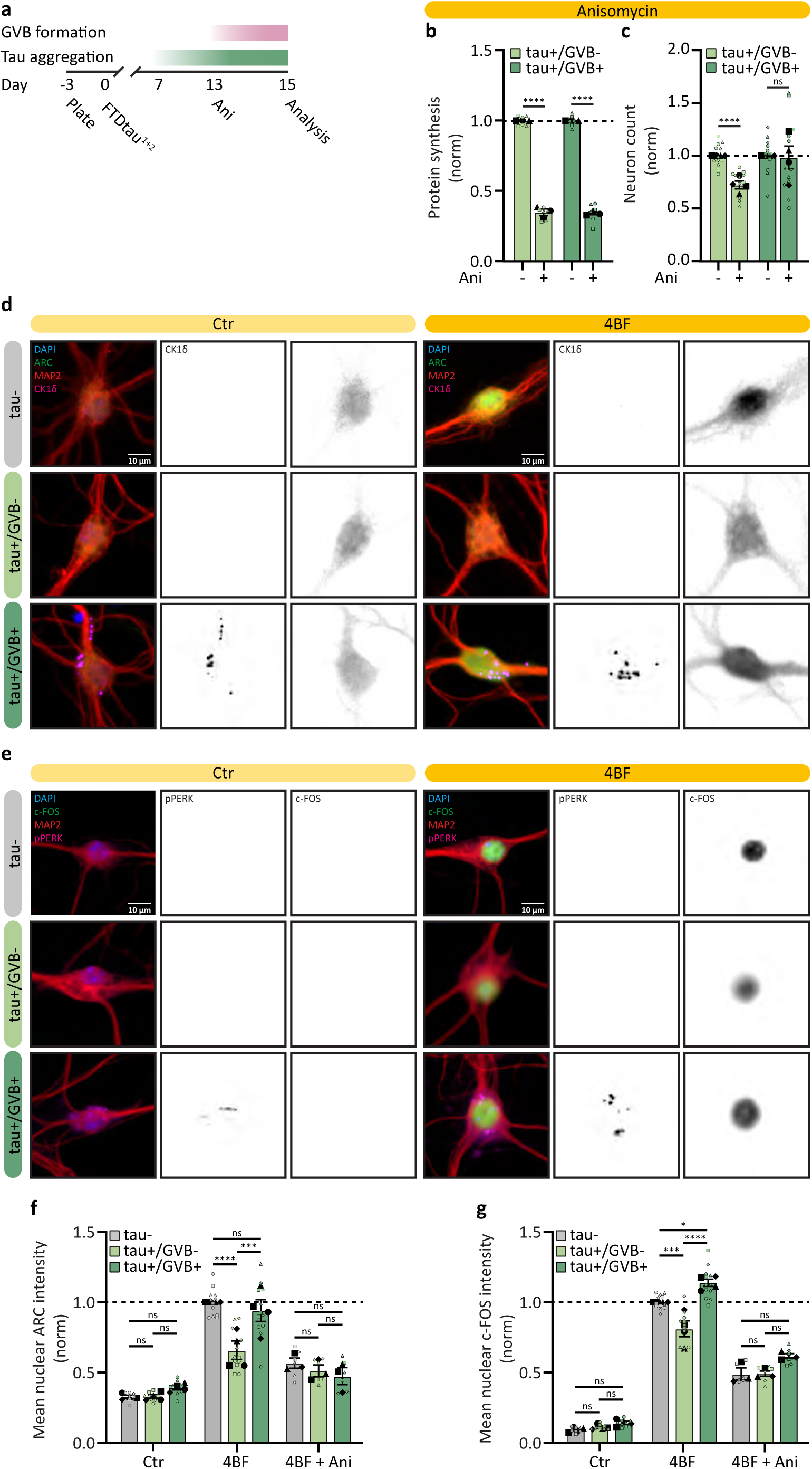
GVB+ neurons fully retain the capacity to control LTP-regulated protein synthesis in the presence of tau pathology. **A** Schematic of the timeline of protein synthesis inhibition using anisomycin (Ani). **B** Quantification of puromycin intensity as a measure for *de novo* protein synthesis in tau+/GVB- and tau+/GVB+ (pPERK based) neurons treated with Ani for 48h analysed at day 15 and normalised to untreated per group. N=3 and n=6. A nested t-test was used. **C** Quantification of neuron number of tau+/GVB- and tau+/GVB+ (pPERK based) neurons treated with Ani for 48h analysed at day 15 and normalised to untreated per group. N=4 and n=12. A nested t-test was used. **D, E** Representative high-content microscopy images of tau-, tau+/GVB- and tau+/GVB+ neurons treated without or with 4BF (see Methods) for 4h. Neurons were immunostained for IEGs ARC (**D**) or c-FOS (**E**), the dendrite marker MAP2 (red) and the GVB marker CK1δ (magenta). Nuclei were visualised by DAPI (blue). Separate channels are shown in greyscales. **F, G** Quantification of mean nuclear ARC (**f**) or c-FOS (**g**) in tau-, tau+/GVB- and tau+/GVB+ (CK1δ (**F**) or pPERK (**G**) based) neurons upon 4BF-dependent LTP-induction treated without or with Ani before LTP-induction, normalised to tau-LTP-induced response. N=4 (Ctr and 4BF) and 3 (4BF + Ani) and n=8, 8 and 8 (Ctr), 12, 12 and 12 (4BF), 6, 6 and 6 (4BF + Ani), for tau-, tau+/GVB- and tau+/GVB+, respectively. A nested one-way ANOVA test followed by a post-hoc Sidak’s test was used. Data are presented as mean ± SEM. *p*-values are indicated: \**p* < 0.05, \*\*\**p* < 0.001, \*\*\*\**p* < 0.0001 and ns: not significant.

To study the relevance of the resilience of GVB+ neurons for neuronal function, we assessed the capacity of tau+ neurons to regulate the acute synthesis of immediate early genes (IEGs) that are involved in the changes of synaptic strength, which is essential for learning and memory-related processes^30,31^. To study if tau aggregation also reduces long-term potentiation (LTP)-related IEG induction in our model, neuronal plasticity was induced using 4BF for 4h after 15 days of tau exposure. 4BF induces chemical LTP and induces the expression of the IEGs ARC and c-FOS^32^. Quantitative high-content automated microscopy was employed to determine expression levels of ARC and c-FOS in the nucleus of tau-, tau+/GVB- and tau+/GVB+ neurons. Both ARC and c-FOS expression were strongly increased upon the induction of chemical LTP (Figures 6D-6G). In tau+/GVB-neurons, ARC and c-FOS levels upon LTP induction were significantly lower than in tau-neurons (decreased by 34% and 19%, respectively), while the tau+/GVB+ neurons do not differ or are even higher in ARC and c-FOS expression levels from tau-neurons, respectively (Figures 6F and 6G). As expected, the upregulation of ARC and c-FOS upon LTP induction was dependent on *de novo* protein synthesis as anisomycin reduced the increase of these proteins (Figures 6F, 6G, S10A and S10B). These data show that GVB+ neurons are resilient to tau-induced impairment in LTP-regulated protein synthesis.

## Discussion

In response to pathological protein assemblies, including tau and α-synuclein, neurons can develop GVBs^14,15^. In the human AD brain -like in our model-the surviving neurons during disease progression are increasingly GVB+^12,33,34^, in accordance with a protective rather than a degenerative role. Our findings demonstrate that CK1δ activity is essential for GVB formation. In addition, we show that autophagy is required for the sequestration of CK1δ into the GVB. The combined data support a model where GVB+ neurons have adapted to tau-induced reduction of protein synthesis, with GVBs as a marker of resilient tau-positive neurons.

Disturbed translational regulation and overall protein synthesis impairment are observed in models of tauopathies^2–6^ and several other neurodegenerative proteinopathies^35–40^. Therefore, impaired protein synthesis is considered an important pathogenic mechanism in neurodegenerative diseases, supported by the amelioration of neurodegenerative phenotypes by the rescue of protein synthesis^5,40–43^. Prolonged reduction of protein synthesis rates affects neuronal viability and function by impairing replacement of critical structural proteins, but also by directly affecting synaptic plasticity, which requires adequately regulated *de novo* protein synthesis^44,45^. The acute and dynamic demand for protein synthesis that is required for synaptic plasticity includes the rapid induction of IEG proteins. Tau pathology has been demonstrated to impair synaptic plasticity, as well as the expression of IEGs in animal models^46,47^. Also in our model, tau aggregation reduces global protein synthesis rates as well as the LTP-regulated induction of IEGs c-FOS and ARC. c-FOS is synthesized in response to increased neuronal activity^48,49^ and ARC is required for the consolidation of LTP and maintenance of synaptic strength^50,51^, both essential for learning and memory-related processes^30,31^. Our data demonstrate that GVB+ neurons not only retain global protein synthesis rates similar to those of neurons without tau pathology, but also importantly retain the capacity to adequately regulate LTP-related IEG protein expression. This demonstrates that GVB+ neurons are resilient to tau pathology-induced protein synthesis decline.

To identify the mechanism by which GVB+ neurons adapt to the tau-induced protein synthesis decrease, transient proteostatic responses mediated by the ISR, mTOR and Angiogenin were investigated. No evidence for their involvement in tau-induced reduction of protein synthesis or the difference between GVB-and GVB+ neurons was found, suggesting that the rescue of protein synthesis involves a more long-term adaptation of the protein synthesis machinery. In AD, progressive ribosomal loss correlates with tau pathology^52^, aligning with studies in tauopathy mouse models that demonstrate decreased ribosomal protein synthesis accompanied by reduced overall protein synthesis^6^. Moreover, targeted laser capture proteomics of human AD hippocampus demonstrated that tangle-bearing neurons exhibit a loss of ribosomal proteins; however, in GVB+ neurons the ribosomal protein levels are similar to those in neurons without tau pathology^53^. We demonstrate here that GVB+ neurons have increased levels of the small ribosomal subunit protein RPS12, the large ribosomal subunit proteins RPL15 and RPL24 as well as the ribosomal biogenesis factors LTV1 and BLTV. Interestingly, in cell lines, CK1δ was previously demonstrated as a critical regulator of ribosome assembly through phosphorylation of LTV1 and BYSL^54–56^. The role of CK1δ in ribosomal biogenesis in neurons is unknown and is a potential candidate for further investigation of GVB-resilience in future studies. This shows that GVB+ neurons have multiple adaptations in the protein biosynthesis machinery that would allow them to counteract the tau pathology-induced decline in protein synthesis. Prolonged protein synthesis impairment by tau pathology will limit the ability to survive additional challenges to the protein synthesis machinery. In line with this, we observe synthetic lethality of pharmacological protein synthesis inhibition and tau pathology in GVB-but not GVB+ neurons. The overall increased protein biosynthesis capacity in GVB+ neurons will result in a healthier proteome that is capable to combat the toxicity of short-term protein synthesis block in GVB+ neurons.

Although GVBs are lysosomes, they do not significantly contribute to the degradation of pathological tau aggregates. Tau does not accumulate within GVBs and the extent of tau pathology is not different between GVB- and GVB+ neurons^15^. GVBs possess degradative capacity similar to the non-GVB lysosomes in the same cell, but are uniquely characterised by the presence of one or multiple dense cores^14^. Live-cell imaging and half-life calculations show that these GVB cores are highly stable. The accumulation of proteins in the GVB core depends on the autophagic machinery, but inhibition of autophagy does not result in accumulation of GVB cores in the cytosol, suggesting that the cores either form within autolysosomes or originate in the cytosol but are rapidly degraded if not engulfed. The basal autophagic machinery is sufficient for the formation of GVBs. CK1δ has been implicated in the early steps of autophagy^57–60^, potentially explaining why CK1δ overexpression can rescue the partial reduction of GVB formation by 48h of Sar405 treatment. Although CK1δ activity is required to form GVB+ neurons, CK1δ activity *in cis* is not required for its localisation to GVBs. These combined data support a model where the accumulation of CK1δ in GVBs occurs by autophagy during the formation of GVB+ neurons. Previous studies showed that CK1δ activity is tightly controlled to prevent dysregulation of the enzyme^61,62^. The strongly increased levels of CK1δ upon inhibition of its activity we observe, suggests that neurons also require tight regulation of CK1δ. Therefore it is not unexpected that there is a direct coupling between the function of CK1δ in initiating the neuronal resilience to tau pathology in GVB+ neurons and the prevention of persisting or excessive activation by sequestering CK1δ in GVBs. The prominent presence of CK1δ in the core could indicate a negative feedback mechanism to sequester the key regulator of the protective mechanism in GVB+ neurons to prevent overactivation of the pathway.

Follow-up studies should address how tau aggregates induce CK1δ activity and what substrates are involved in resilience. The number of neurons with tau pathology that become GVB+ increases over time, suggesting a (partially) stochastic process. CK1δ is traditionally considered a constitutively active protein, relying on binding partners or scaffold proteins for subcellular localisation^61,63^, although also differential autophosphorylation was more recently shown to regulate substrate selectivity^64^. CK1δ binds to and phosphorylates tau at different sites^61^. Tau assemblies may provide a platform for CK1δ binding and signalling, leading to GVB formation. This may involve structural rather than primary sequence motifs because GVBs also form in response to α-synuclein aggregates^15^, although the involvement of CK1δ and protein synthesis resilience in α-synuclein-induced GVBs has currently not been demonstrated.

The identification of specific molecular targets to stimulate GVB formation has therapeutic potential. Because GVBs form selectively in neurons, it is likely that, in addition to CK1δ, cell type-specific factors are involved. Ultimately, combining resilience-boosting therapy with tau-reducing therapies like *MAPT* antisense oligonucleotides or tau immunotherapy is an attractive scenario to halt the disease while also protecting neurons against the cellular effects of tau pathology to help them to retain function. By elucidating the mechanistic underpinnings of the formation and functional implications of the presence of GVBs in neurons, our work offers new insights into neuronal resilience in tauopathies.

## Limitations of the study

CK1δ inhibition can completely block the generation of GVBs and thereby the resilient GVB+ neurons. The increased levels of ribosomal proteins assembly factors in GVB+ neurons together with lack of evidence for transient stress response regulation support a mechanism involving ribosomal biogenesis in the resilience. Currently, the lack of interventions that can disconnect the resilience program in GVB+ neurons and the formation of GVBs prevents straightforward investigation of the mechanistic relation between CK1δ and the protein synthesis phenotypes. This also complicates a more definite conclusion that GVBs are merely a byproduct of activation of the protective mechanism in GVB+ neurons. However, currently, there is no indication for a specific function associated with GVBs themselves.

## Resource availability

### Lead contact

Requests for further information and resources should be directed to the lead contact, Wiep Scheper.

## Materials availability

All unique/stable reagents generated in this study are available upon reasonable request from the lead contact with a completed materials transfer agreement (MTA).

## Data and code availability

- The mass spectrometry proteomics data have been deposited to the ProteomeXchange Consortium via the PRIDE^65^ partner repository with the dataset identifier PXD061887.
- This paper does not report original code.
- Any additional information required to reanalyse the data reported in this work paper is available from the lead contact upon reasonable request.

## Acknowledgements

We thank Bart Klein for expert technical support, Joke Wortel for animal breeding and Robbert Zalm and Ingrid Saarloos for plasmid construction and lentivirus production. We also would like to thank the Molecular Neurodegeneration group and our Functional Genomics Department colleagues for stimulating discussions and critical reading of the manuscript. This work was supported by Hersenstichting/Coby van Nieuwkerkfonds #DR-2019-00278, Alzheimer Nederland #WE.03-2017-10 and The Netherlands Organisation for Scientific Research (NWO) #OCENW.M.21.123 to WS.

## Author contributions

J.F.M.S., T.W.L., M.J.O., S.M. and F.S. performed the experiments and analysed the results. D.P.I. and K.W.L. performed the LC-MS/MS. Writing of the original draft was done by J.F.M.S. and W.S. Reviewing was done by all authors. J.F.M.S. and W.S. conceived the study and W.S. provided supervision and obtained funding for the study.

## Methods

### Animals and primary cell culture

Animals were kept and bred according to institutional and Dutch governmental guidelines and experiments were approved by the animal ethical committee of the VU University/VU University Medical Center. C57BL/6 mice were bred in-house or obtained from Charles River.

Embryonic day (E) 18.5 WT mouse embryos or postnatal day 1 (P1) mice were dissected to obtain cerebral cortices after disposal of the meninges. The cortices were digested in Hanks balanced salt solution (HBSS, Sigma-Aldrich) containing 10 mM HEPES (Gibco; Hanks-HEPES) and 0.25% trypsin (Gibco) for 20 min at 37°C. Hereafter, the digested tissue was washed twice with Hanks-HEPES and subsequently with DMEM (Lonza) containing 10% heat-inactivated fetal bovine serum (Gibco), 1% penicillin-streptomycin (Gibco) and 1% non-essential amino acid solution (Gibco; DMEM+) in which the tissue was triturated using a fire-polished Pasteur pipette. Subsequently, the dissociated cells were spun down for 5 min at 800g at room temperature (RT), resuspended and plated in neurobasal medium (Gibco) supplemented with 2% B-27 (Gibco), 18 mM HEPES, 0.25% Glutamax (Gibco) and 0.1% penicillin-streptomycin (NB+).

Primary cortical neurons were cultured in plates and glass coverslips coated with 5 µg/mL poly-L-ornithine (Sigma-Aldrich) and 2.5 μg/mL laminin (Sigma-Aldrich and Bio-Techne) overnight (O/N) at room temperature (RT). All cells were maintained at 37°C and 5% CO_2_. For Western blotting, cortical neurons were plated in a 6-well plate at a density of 300.000 neurons per well. For immunolabeling, neurons were plated on 13 mm glass coverslips or in a 96-well plate (Greiner) at a density of 40.000 cells or 12.500 cells per well, respectively. After 10 days *in vitro* (DIV10), fresh NB+ was supplied to each well at 40% of the total volume in the well.

### Lentiviral transduction in primary cells

Lentiviral transduction with the human 2N4R P301L, S320F tau (FTDtau^1+2^) and FTDtau^1+2^ with an in-frame C-terminal EGFP tag (FTDtau^1+2^-GFP) were performed on DIV3 (Day0), unless otherwise specified, in primary neurons^15,16,23^.

WT mouse CK1δ (kind gift from Dr. Y.E. Greer) has been described before^66^ and the formation of N-terminally tagged CK1δ with EGFP (GFP-CK1δ) has been described before^14^. GFP-CK1δ_RES_ was changed from WT CK1δ in the *shCsnk1d* target sequence to prevent KD (Table 1) (new sequence: CGGGA**C**CG**T**GA**G**GAACGATTA). A nuclear export signal (NES) was N-terminally cloned into the GFP-CK1δ construct. For the kinase-dead variant of CK1δ, a single point-mutation in lysine-38 to methionine-38 (AAG to ATG) was created to form GFP-CK1δ-K38M. As a control for EGFP-tagged constructs, a lentiviral CMV-driven EGFP construct was used. All constructs were subcloned into lentiviral backbone vectors under the cytomegalovirus (CMV), the neuron-specific synapsin (Syn) or a doxycycline-inducible Syn promoter using the Gateway system (Invitrogen). The production of lentiviral particles was performed as described previously^67^ and the lentiviral particles were stored at -80°C until use.

**Table 1:**
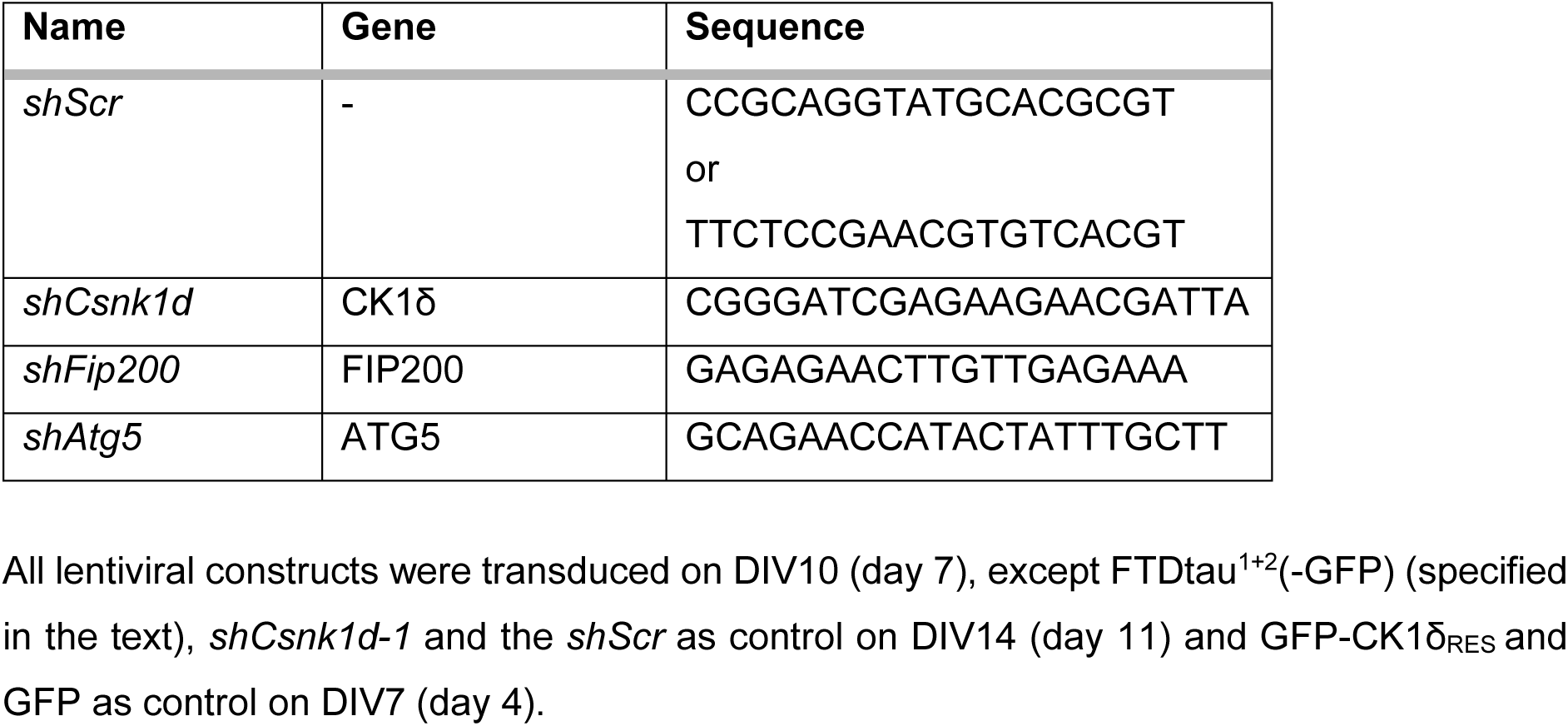
The names, targeting gene and sequence of the shRNA of all shRNAs used in this research.

Plasmids for short hairpin RNA (shRNA) constructs for the scrambled control (*shScr*) and CK1δ (Sigma-Aldrich; TRCN0000023773) were obtained from the MISSION shRNA library (Sigma-Aldrich). shRNAs targeting FIP200 (kind gift of Dr. Marijn Kuijpers)^26^ and ATG5 were cloned using oligonucleotides (Sigma-Aldrich) based on previously published research^27^. The shRNAs were cloned into a second-generation lentiviral backbone under a CMV promoter.

All lentiviral constructs were transduced on DIV10 (day 7), except FTDtau^1+2^(-GFP) (specified in the text), *shCsnk1d-1* and the *shScr* as control on DIV14 (day 11) and GFP-CK1δ_RES_ and GFP as control on DIV7 (day 4).

### Cell treatments

Primary neurons were treated with 50 μM E64D and 20 μM Pepstatin-A (Sigma-Aldrich) for 24h. 5 μM of Sar405 (Sigma-Aldrich) was used for indicated times to inhibit VPS34 and 5 μM Torin1 (Bioke) (72h, 48h or 24h) or 1 μM rapamycin (Hello Bio) (48h) were used to inhibit mTOR. To inhibit CK1δ, neurons were treated with PF-670462 (CK1δ_i_, TargetMol) for indicated times at a concentration of 12.5 µM. To induce ER-stress, primary neurons were treated with tunicamycin (TM, Sigma-Aldrich) for 24h at 5 µg/ml. 200 nM of the ISR inhibitor ISRIB (Selleck Chemicals) was added 2h before fixation. Angiogenin inhibitor (NSC-65828, National Cancer Institute) was added 24h before fixation at a final concentration of 75 mM and was previously validated^29^. Anisomycin (Sigma-Aldrich) was added for 48h, or directly before the induction of LTP, at a final concentration of 50 ng/ml to determine the effect on global protein synthesis. To label *de novo* synthesised proteins, we performed surface sensing of translation (SUnSET) as previously described^28^. In short, primary cells were incubated with 2 µM puromycin (InvivoGen) for 15 min before fixation. Using the anti-puromycin antibody (see Table 2), puromycinylated proteins were detected. To stain nucleic acids, cells were incubated with 1 µM SYTO RNASelect (Thermo Fisher Scientific) for 4 hr before methanol fixation. Compounds were dissolved in H_2_O (puromycin) or DMSO (the rest) and diluted in NB+. Equal amounts of solvent were added to the control conditions or when multiple concentrations were used, the highest concentration was used to determine the amount of vehicle added.

**Table 2:**
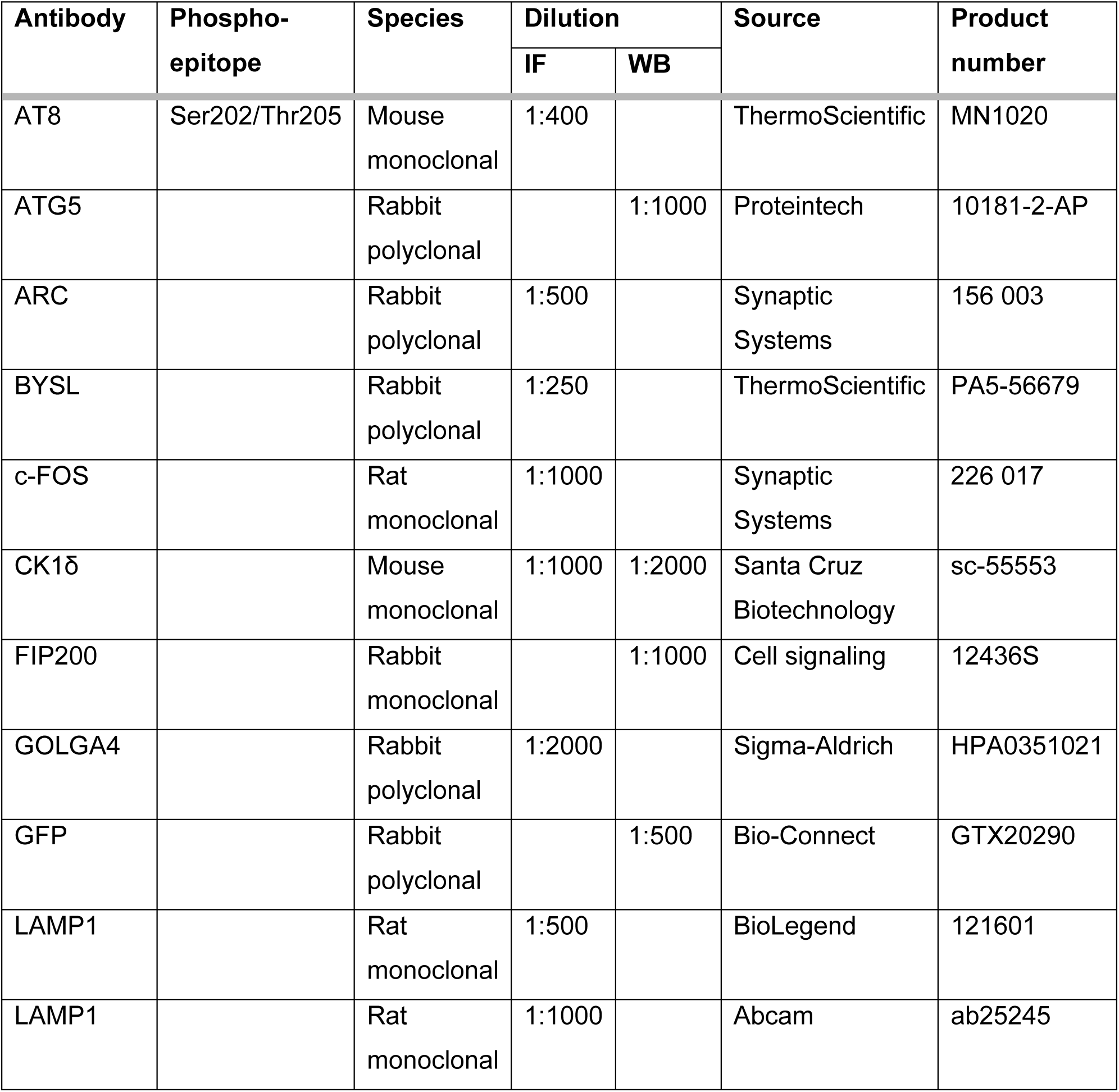
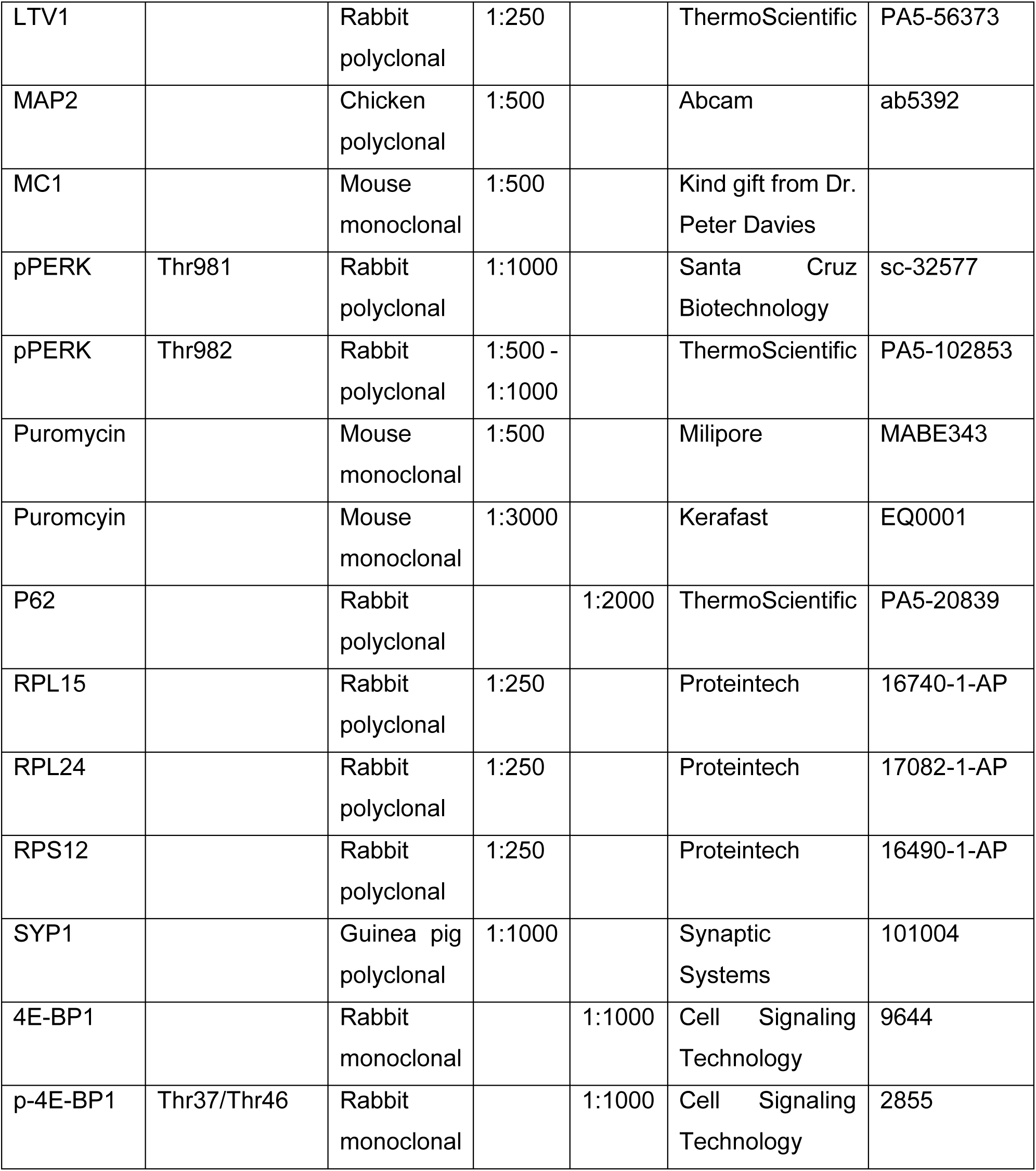
Antibodies used in this research. From left to right: The target protein the primary antibody was raised at; for antibodies targeting specific phosphor-sites, the phospho-epitope is listed; the species of the primary antibody; the dilution used for immunofluorescence (IF) or western blot (WB); the source; the product number.

| Antibody | Phospho-epitope | Species | Dilution |  | Source | Product number |
| --- | --- | --- | --- | --- | --- | --- |
|  |  |  | IF | WB |  |  |
| AT8 | Ser202/Thr205 | Mouse monoclonal | 1:400 |  | ThermoScientific | MN1020 |
| ATG5 |  | Rabbit polyclonal |  | 1:1000 | Proteintech | 10181-2-AP |
| ARC |  | Rabbit polyclonal | 1:500 |  | Synaptic Systems | 156 003 |
| BYSL |  | Rabbit polyclonal | 1:250 |  | ThermoScientific | PA5-56679 |
| c-FOS |  | Rat monoclonal | 1:1000 |  | Synaptic Systems | 226 017 |
| CK1δ |  | Mouse monoclonal | 1:1000 | 1:2000 | Santa Cruz Biotechnology | sc-55553 |
| FIP200 |  | Rabbit monoclonal |  | 1:1000 | Cell signaling | 12436S |
| GOLGA4 |  | Rabbit polyclonal | 1:2000 |  | Sigma-Aldrich | HPA0351021 |
| GFP |  | Rabbit polyclonal |  | 1:500 | Bio-Connect | GTX20290 |
| LAMP1 |  | Rat monoclonal | 1:500 |  | BioLegend | 121601 |
| LAMP1 |  | Rat monoclonal | 1:1000 |  | Abcam | ab25245 |
| LTV1 |  | Rabbit polyclonal | 1:250 |  | ThermoScientific | PA5-56373 |
| MAP2 |  | Chicken polyclonal | 1:500 |  | Abcam | ab5392 |
| MC1 |  | Mouse monoclonal | 1:500 |  | Kind gift from Dr. Peter Davies |  |
| pPERK | Thr981 | Rabbit polyclonal | 1:1000 |  | Santa Cruz Biotechnology | sc-32577 |
| pPERK | Thr982 | Rabbit polyclonal | 1:500 - 1:1000 |  | ThermoScientific | PA5-102853 |
| Puromycin |  | Mouse monoclonal | 1:500 |  | Milipore | MABE343 |
| Puromycin |  | Mouse monoclonal | 1:3000 |  | Kerafast | EQ0001 |
| P62 |  | Rabbit polyclonal |  | 1:2000 | ThermoScientific | PA5-20839 |
| RPL15 |  | Rabbit polyclonal | 1:250 |  | Proteintech | 16740-1-AP |
| RPL24 |  | Rabbit polyclonal | 1:250 |  | Proteintech | 17082-1-AP |
| RPS12 |  | Rabbit polyclonal | 1:250 |  | Proteintech | 16490-1-AP |
| SYN1 |  | Guinea pig polyclonal | 1:1000 |  | Synaptic Systems | 101004 |
| 4E-BP1 |  | Rabbit monoclonal |  | 1:1000 | Cell Signaling Technology | 9644 |
| p-4E-BP1 | Thr37/Thr46 | Rabbit monoclonal |  | 1:1000 | Cell Signaling Technology | 2855 |

### Induction of chemical LTP

Neurons at DIV18 (day 15) were stimulated for 4 hours, after which they were directly fixed. A cocktail of chemicals was used to induce network activity and plasticity for generation of immediate early gene (IEG) expression. The 4BF cocktail consists of 100 µM 4-aminopyridine (Sigma, A78403), 50 µM bicuculline (Sigma, 14343) and 50 µM forskolin (Sigma, 344282) as previously described^32^.

### Fixation and immunolabeling of primary neurons

On DIV18 (day 15), DIV25 (day 22) or DIV32 (day 29) neurons were fixed in 1.85% formaldehyde (FA, Electron Microscopy Sciences) in PBS (pH 7.4) by the addition of equal volumes of 3.7% FA to the culture medium for 10 min followed by fixation in 3.7% FA for 10 min at RT. Primary neurons shown in Figure 5M and Figure S4V were fixed in ice-cold methanol for 15 min on ice to remove soluble proteins. After fixation, cells were washed with PBS and stored at 4°C in PBS (pH 7.4) in the dark until further use. Fixed cells were permeabilised in 0.5% Triton X-100 (Thermo Fisher Scientific) in PBS (pH 7.4) for 5 min at RT and blocked in 2% normal goat serum (Gibco) and 0.1% Triton X-100 in PBS (blocking solution) for 30 min at RT. Primary antibodies were diluted in blocking solution and incubated O/N at 4°C in the dark. The primary antibodies used and their dilutions are listed in Table 2. After 3 PBS washes, cells were incubated with Alexa Fluor (405, 488, 546, 568, 647)-conjugated secondary antibodies (Abcam and Thermo Fisher Scientific) in a 1:500 dilution in blocking solution for 1 h at RT in the dark. Cells were subsequently washed thrice with PBS. Aggresomes were visualised using the proteostat aggresome detection kit (Enzo) according to the manufacturer’s details and washed three times with PBS. Hereafter, to visualise nuclei, cells were incubated with DAPI (Brunschwig Chemie) diluted in PBS (5 μg/mL) at RT for 5 min, followed by two PBS washes. Coverslips were mounted on microscope slides (Thermo Fisher Scientific) using Mowiol (Sigma-Aldrich) and were kept in the dark O/N at RT to dry. Slides were stored in the dark at 4°C until imaging. 96-well plates were stored in the dark at 4°C until imaging with the wells filled with 100 μl PBS.

### Confocal microscopy and confocal image analysis

The mounted coverslips were imaged using a Nikon Eclipse Ti confocal microscope controlled by NisElements 4.30 software (Nikon), equipped with a 60x oil immersion objective (NA = 1.4) or a 40x oil immersion objective (NA = 1.3). Z-stacks with a step size of 0.25, 0.5 or 1 μm were obtained. Laser settings were adjusted to reduce saturation and kept the same in every independent experiment between conditions.

### Live microscopy

The cells were placed in a Tokai Hit live-cell microscope incubation system mounted in the Nikon Eclipse Ti confocal microscope, equipped with a 40x oil immersion objective (NA = 1.3) or a 10x air objective (NA = 0.45) for calcium imaging. 10 min before calcium imaging, 0.5 μM X-Rhod-1 (Thermo Fisher Scientific) was added to the cells. A video with 8fps was made for the duration of 5 min. For following GVB dynamics, every 10, 20 or 30 min for a period of minimally 9h, a Z-stack with a step size of 1 μm was obtained. When FRAP was applied, a specified region in the GVB+ soma was photobleached and followed over the same period as described in the text.

Confocal images were analysed using ImageJ software (National Institutes of Health). Neurons with 2 or more CK1δ or pPERK bright punctae were defined as GVB+ neurons and neurons without any punctae were defined as GVB-neurons. Single confocal planes are shown unless otherwise stated.

Line segments were drawn to determine the overlap between multiple markers. The line graphs show fluorescence intensity in arbitrary units (AU).

### High-throughput microscopy and automated analysis

Cells cultured in a 96-well plate were imaged using CellInsight CX7 High-Content Screening (HCS) Platform (Thermo Fisher Scientific) controlled by HCS Studio Cell Analysis software (Thermo Fisher Scientific). 10x, 20x and 40x air objectives were used in widefield mode. The 10x or 20x objectives were used to determine toxicity of the treatment compared to control. The 20x objective was used for determination of morphology. The 20x and 40x objectives were used to measure intensity and determine the number of GVB+ neurons. A single focal plane was obtained for all channels. 25 or 30 fields, distributed throughout the well, were imaged for 10x or 20x and 40x, respectively. In all cases, DAPI signal was used for autofocus for every image individually. On average, 10x imaging obtained data from 1000-10000 neurons per well, 20x imaging from 50-1000 neurons per well and 40x 25-500 neurons per well.

The data obtained from the high-throughput microscopy were analysed with Columbus analysis software (v2.5.2.124862; PerkinElmer) by in-house developed scripts. Each script was optimized per independent experiment. In every analysis, neuronal nuclei were defined by overlap between the nuclear marker DAPI and the neuron-specific marker MAP2 as well as the size and roundness of the nucleus. Outcome parameters were averaged per well (when applicable) and normalised to the mean of the control wells. These values were used in the analysis.

### Neuronal morphology

Dendritic morphology and pre-synaptic density were determined using in-house developed scripts, previously published^23,29^. In short, based on MAP2 staining, dendritic morphology was determined using a built-in CSIRO neurite analysis module. Morphological features, including dendrite length, number of extremities, segments and primary dendrites were normalised to the number of neuronal nuclei within that well. The amount of synaptophysin 1 (SYP1) positive pre-synapses was determined based on local intensity maxima within an enlarged MAP2 mask. The pre-synapse density was determined by dividing the total amount or SYP1 puncta over total dendrite length.

### Tau intensity in MAP2 mask

The intensity measurements of multiple readouts of tau aggregation, GFP-tagged and untagged FTDTau^1+2^, were performed by measuring the sum of the intensity of FTDTau^1+2^-GFP or the conformational pathogenic tau marker MC1 in a MAP2 mask. The absolute intensity was normalised to the total area of the MAP2 mask.

### GVB selection

GVB+ neurons were defined using an altered protocol, as previously described^14,15^. In short, the GVB marker (pPERK, CK1δ, Dox-GFP-CK1δ or GolginA4) was subjected to a filter step to decrease background intensity using a curvature optimized per marker. Single puncta were selected based on intensity. The single puncta were subsequently clustered based on their proximity to one another. These clusters were defined as GVB-clusters based on their size, roundness, intensity of the GVB-marker, the intensity of MAP2 in the cluster and exclusion of the cluster within the nucleus based on DAPI-signal. The percentage of GVB-clusters was calculated by dividing the number of GVB-clusters by the number of neurons in the same well.

### Somatic and nuclear intensity

The soma was defined as the MAP2-positive signal 13µm around the neuronal nucleus (excluding overlap between bordering somas), after smoothing and filling the holes of the MAP2 signal. Subsequently, somas in neurons transduced with FTDtau^1+2^ were defined as GVB-or GVB+ as previously described. False positives were excluded in tau-neurons. Within each soma, the mean intensity of the marker (e.g. puromycin, RPS12, RPL15, RPL24, LTV1 and BYSL) was measured in the complete soma (cytosol and nucleus). Somatic CK1δ levels were determined by measuring CK1δ intensity in the soma without the GVBs. Nuclear intensity for CK1δ was corrected by subtracting the mean nuclear intensity by the mean intensity level of the cytosol without the nucleus. When determining CK1δ levels in a GVB+ neuron, pPERK was used to define GVB-positivity. The nuclear intensity of c-FOS and ARC were not background corrected. The number of GVB+ somas was divided by the number of somas to calculate the percentage of GVB+ somas.

### GVB selection in GFP-positive neurons

For the selection of GFP-positive (GFP+) neurons, the soma was defined as previously described. Mean GFP intensity was measured in the complete soma region, including the nucleus. Based on an untransduced negative control, a threshold was set to select neurons with a higher somatic mean GFP-intensity than the negative control. These neurons were defined as GFP+ neurons. GVB-clusters were defined as GFP+ if they were localised within the GFP+ soma, creating GVB-/GFP+ and GVB+/GFP+ somas. Within these two populations, mean GFP-intensity was measured in the complete soma (cytosol and nucleus), only the nucleus and the cytosol without the nucleus. Nuclear intensity was corrected by subtracting the mean nuclear intensity by the mean intensity level of the cytosol excluding the nucleus and GVBs, per neuron. The number of GVB+/GFP+ neurons was divided by the number of GFP+ neurons to calculate the percentage of GVB+/GFP+ neurons.

### Primary neuron lysis and western blot analysis

Cells were washed 2x with ice-cold PBS and samples for the isolation of total proteins were scraped in loading buffer containing dithiothreitol (50mM) and 2% sodium dodecyl sulfate (SDS) to lyse the cells. The samples were triturated by a 21-gauge needle to sheer the DNA. For analysis of cytosolic proteins, cells were lysed in 1%-Triton X-100 (Thermo Fisher Scientific) in PBS supplemented with cOmplete protease inhibitor cocktail (Roche) and PhosSTOP phosphatase inhibitors (Roche). To purify these whole-cell lysates, the lysates were centrifuged at 14.000 g for 10 min at 4°C. The total protein content of the supernatant was determined with the Pierce BCA protein assay kit (BCA) protein assay (Thermo Fisher Scientific). All samples were boiled for 5 min at 95°C. An equal volume or amount of the samples was loaded to a 4-15% Mini-Protean TGX stain-free precast polyacrylamide gels (Bio-Rad) and Precision Plus ProteinTM All Blue Standards (Bio-Rad) was used as a ladder. The total amount of protein on the gel was visualised using the Gel Doc EZ System (Bio-Rad) and analysed using Image LabTM 6.0 (Bio-Rad). The total protein content was determined by correcting for the measured area size. The samples were then transferred onto 0.2 μM nitrocellulose membranes with the TransBlot Turbo transfer system and kit (both from Biorad).

The membranes were then blocked with 5% skimmed milk powder (Millipore) in TBS containing 0.05% Tween (TBS-T) (Sigma-Aldrich) for 30 min at RT. Afterwards, the membranes were incubated with primary antibody diluted in blocking buffer overnight at 4°C (See Table 2 for the dilutions of the primary antibodies). Subsequently, the membranes were washed 4x 15 min in TBS-T whereafter the membranes were incubated with HRP-conjugated secondary antibodies (DAKO) diluted in blocking buffer (1:2000) for 1 hour at RT. Last, the membranes were washed 4x 15 min with TBS-T and submerged with SuperSignalTM West Femto Maximum Sensitivity Substrate kit (Thermo Fisher Scientific). Membranes were stripped for reblotting using ReBlot Plus Strong Antibody Stripping Solution (Merck). The chemiluminescence was visualized with the Odessey Fc system (Li-Cor) and analysed using Image Studio 6.0 software (Li-Cor). Band intensities were corrected for the background signal.

## Proteomics analysis of primary neurons

### Sample preparation

At day 15, media was aspirated and neurons were subjected to 2x gentle ice-cold PBS washes. Neurons were then collected into 1.5 mL tubes using ice-cold PBS supplemented with protease inhibitor (Roche). Samples were centrifuged for 5 min at 3000 x g at 4°C and the remaining pellet was resuspended in 7 µL of 5x loading buffer containing 10% SDS, 0.25 M Tris pH 6.8, 0.1% bromophenol blue, 0.5 M DTT, 50% glycerol. Samples were stored at −20°C until further use. A total of 14 samples from two independent experiments were employed per condition.

### Protein in-gel digestion

Cell lysates were incubated at 98°C for 6 min to denature the proteins followed by incubation with 30% acrylamide/Bis Solution 37.5:1 (Bio-Rad) for 30 min at RT to block cysteine residues. Samples were loaded onto 1 mm thick acrylamide gels. They were composed of 10% acrylamide, 0.375 M Tris-HCl (pH 8.8), ultra-pure H_2_O, 0.1% (w/v) APS and 6 µL N,N,N’,N’-tetramethylethylene-diamine (TEMED; Biorad) per gel. Proteins were allowed to migrate into the gel by electrophoresis (120 V) for approximately 10 min. Peptides were extracted as previously described^68^. Briefly, gels were fixed O/N in a solution containing 50% (v/v) ethanol and 3% (v/v) phosphoric acid in H_2_O at RT and stained with Colloidal Coomassie Blue (34% (v/v) methanol, 3% (v/v) phosphoric acid, 15% (w/v) ammonium sulphate and 0.1% (w/v) Coomassie brilliant blue G-250 (Sigma-Aldrich)). Sample-containing lanes were separated and cut in blocks. Gel fragments were destained in 50 mM NH_4_HCO_3_ (99% NH_4_HCO_3_, Fluka) and 50% (v/v) acetonitrile (HPLC grade; JT Baker), dehydrated using 100% acetonitrile, re-swell in 50 mM NH_4_HCO_3_ containing 10 µg/ml trypsin (sequence grade; Promega) and incubated O/N in a humidified chamber at 37°C to facilitate protein digestion. Peptides were extracted with a solution containing 0.1% (v/v) trifluoroacetic acid (Sigma-Aldrich) and 50% (v/v) acetonitrile. Samples were dried using a SpeedVac (Eppendorf), and stored at −20°C until LC-MS analysis.

### LC-MS analysis

Samples (75 ng) were loaded into EvoTips (EV2003; Evosep) and run on a 15 cm x 75 µm, 1.9 µm Performance Column (EV112; Evosep) using the Evosep One LC system with 30 samples per day program. Peptides were electro-sprayed into the TimsTOF Pro 2 mass spectrometer (Bruker Daltonics) and analysed with diaPASEF^69^. The MS was operated with the following settings: scan range 100-1700 *m/z*, ion mobility 0.6 to 1.6 Vs/cm^2^, ramp time 100 ms, accumulation time 100 ms and ramp rate 9.42 Hz.

### Data analysis

Raw diaPASEF data from the TimsTOF Pro 2 were searched with a virtual spectral library generated from the UniProt mouse proteome UP000000589_1090 using DIA-NN 1.8.1^70^. Deep-learning-based spectra and cross-run normalisation were activated. The maximum number of missed cleavages was set to 1. The peptide length range was 7-30, the precursor charge range to 2-4, the precursor m/z range to 300-1800 and the fragment ion m/z range to 200-800. A fixed modification of UniMod: 24, 71.0371 at C was used, which represents acrylamide adduct. The precursor False Discovery Rate (FDR) was 1% (default). The rest of the settings were used as default.

MS-DAP (v. 1.0)^71^ was used for quality control and candidate discovery on the results obtained from DIA-NN. Only peptides identified and quantified in at least 6 samples from each group were included in the analysis. The batch effect of samples from different experiments was removed by including the variable “batch” in MS-DAP. Normalisation of peptide abundance values was applied and the MSqRob algorithm was used for differential analysis at the peptide level using an FDR threshold of 5%.

GO and pathway analysis was performed using ShinyGO (v. 0.81)^72^ with an FDR-adjusted p-value threshold of 0.05. All detected proteins were used as background dataset. The glial genes GFAP, OLIG1, OMG and AQP4 (as shown by squares in Figure 5A) were omitted for the GO-term analysis. A minimum of 3 proteins belonging to the same pathway and a maximum of 500 were considered to identify that pathway as a hit.

The mass spectrometry proteomics data have been deposited to the ProteomeXchange Consortium via the PRIDE^65^ partner repository with the dataset identifier PXD061887.

## Statistics

For each experiment, at least 3 biological replicates were performed, containing multiple technical replicates (as indicated in the figure legends). Biological replicates are indicated in large, black shapes in the figures, while technical replicates belonging to the same biological replicate are smaller, lighter and of the same shape, except in Figure 1E-1G, for visual clarity. Data were normalised to the mean of the control condition, per timepoint unless otherwise specified. GraphPad Prism 8.4.2 software was used for statistical analysis. Visual, technical outliers in the pooled data were tested using the ROUT method (Q = 1%) and excluded when significant. A nested t-test was used for the comparison of two groups or a nested one-way ANOVA followed by either a Dunnett’s or a Sidak’s multiple comparison post-hoc test for the comparison of more than two groups^73^. When only tau+/GVB-and tau+/GVB+ neurons from the same well were compared, a paired t-test was used. For western blot analyses, an unpaired t-test was used to compare two groups or a one-way ANOVA followed by a Sidak’s post-hoc test was used when applicable. Only the most relevant *p*-values were included in the figures. A *p*-value of < 0.05 was considered statistically significant. \**p* < 0.05, \*\**p* < 0.01, \*\*\**p* < 0.001, \*\*\*\**p* < 0.0001 and ns.

## Declaration of generative AI and AI-assisted technologies in the writing process

During the preparation of this work, we used ChatGPT in order to restructure individual sentences. After using this tool/service, we reviewed and edited the content as needed and take full responsibility for the content of the publication.

---

**Figure S1:**
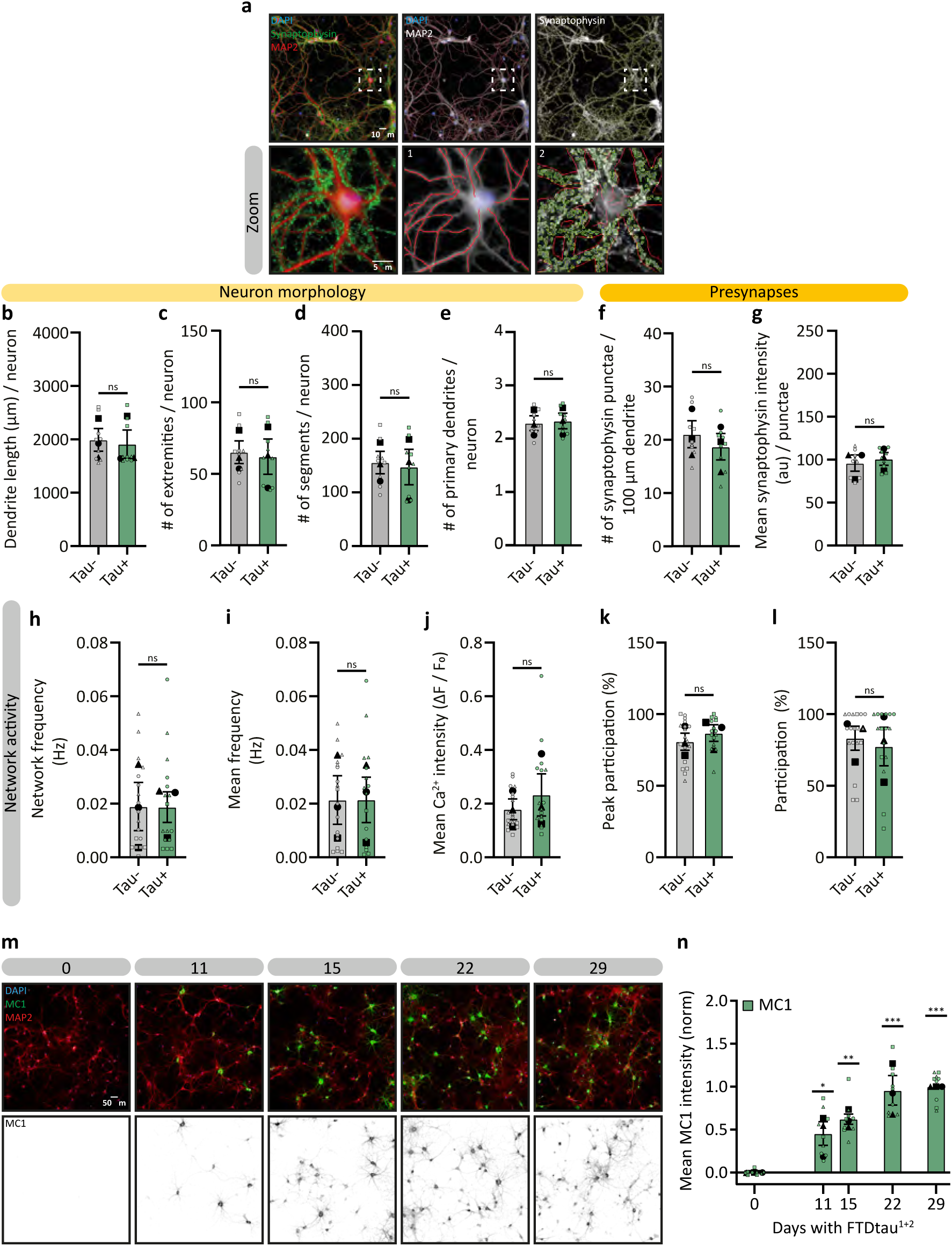
Tau aggregation does not alter neuronal morphology, viability and network activity, related to Figure 1. **A** Representative high-content microscopy images showing the workflow for assessing neuron and pre-synapse morphology based on immunofluorescence staining of the dendrite marker MAP2 (red, grey) and the pre-synapse marker synaptophysin (green, grey) analysed at day 15. Nuclei were visualised by DAPI (blue). The zoomed regions are indicated by dashed white squares. The detection of dendrites is based on the MAP2 signal (red traces, 1) and pre-synapses (yellow outline, 2) are defined using synaptophysin punctae within the dendrite region (red outline, 2). **B-G** Quantification of neuron and pre-synapse morphology in tau- or tau+ neurons analysed at day 15, including dendrite length (**B**), number of extremities (**C**), number of segments (**D**), and number of primary dendrites (**E**), normalised to neuron number. Pre-synaptic analysis includes pre-synapse density (**F**) and synaptophysin intensity (**G**) normalised to dendrite length. N=3 and n=9. A nested t-test was used. **H-L**: Quantification of network activity in tau-or tau+ neuronal networks using calcium imaging analysed at day 15 including network frequency (**H**), mean frequency (**i**), mean calcium intensity (**J**), , peak participation (**K**) and participation (**L**). Single datapoints represent recordings from 10 neurons per field of view. N=3 and n=18 (tau-) or 16 (tau+). A nested t-test was used. **M** Representative high-content microscopy images at different FTDtau^1+2^ exposure durations as indicated. Neurons were immunostained for the pathological tau marker MC1 (green) and MAP2 (red) and nuclei were visualised by DAPI (blue). MC1 channel is shown in greyscale. **N** Quantification of mean MC1 immunofluorescence intensity normalised to 29 days FTDtau^1+2^ exposure (1) and untransduced (0). N=3 and n=9 for all FTDtau^1+2^-exposures. A nested one-way ANOVA followed by a Dunnett’s post-hoc test was used. Data are presented as mean ± SEM. *p*-values are indicated: \**p* < 0.05, \*\**p* < 0.01, \*\*\**p* < 0.001 and ns: not significant.

**Figure S2:**
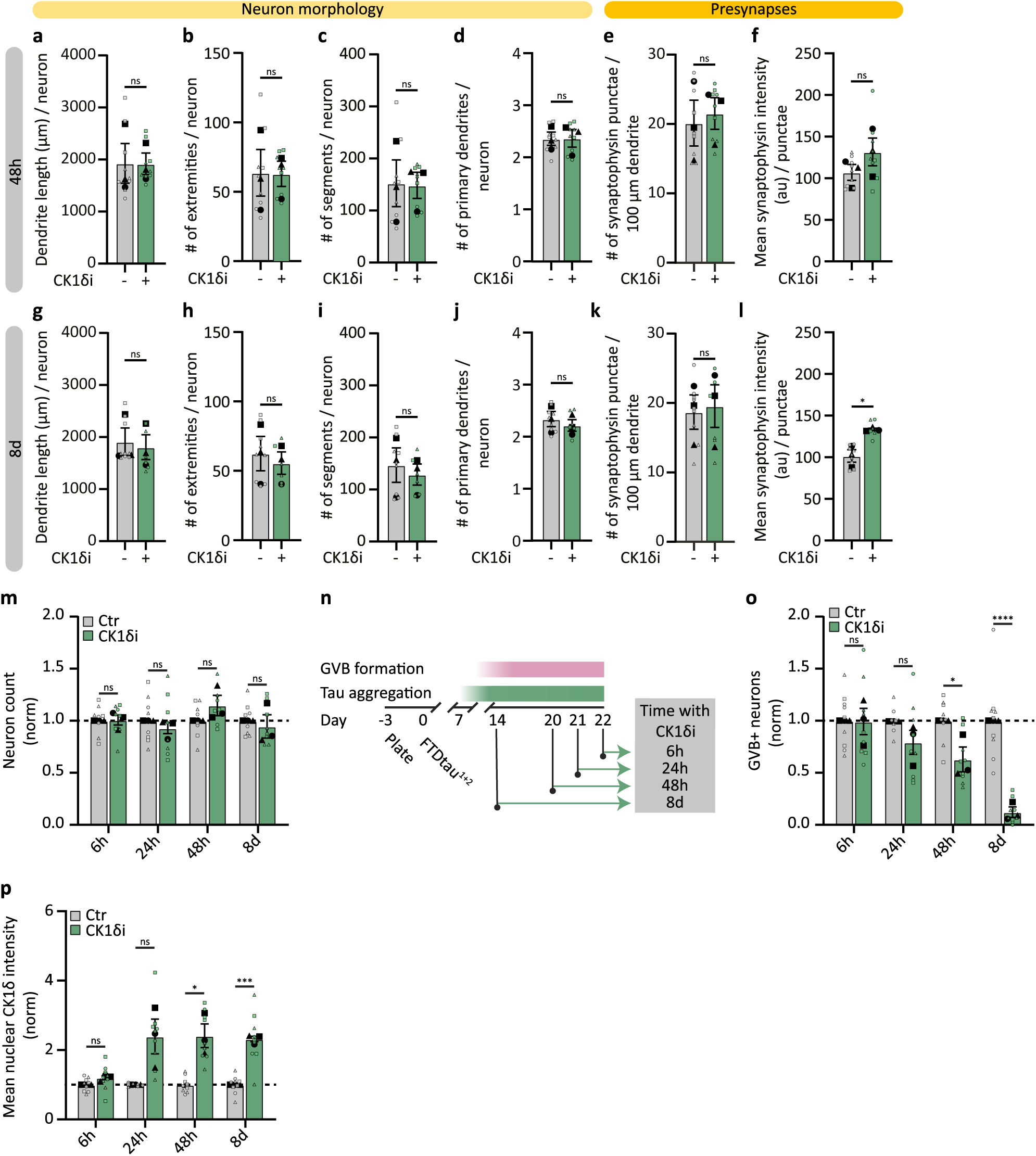
CK1δ inhibition not does alter neuronal morphology, related to Figure 2. **A-l** Quantification of neuron and pre-synapse morphology of tau+ neurons treated without or with CK1δ_i_ for 48h or 8d analysed at day 15. Dendrite length (**A, G**), number of extremities (**B, H**), number of segments (**C, I**) and number of primary dendrites (**D, J** were normalised to number of neurons. Number of pre-synapses (**E, K**) was normalised to dendrite length and the intensity of synaptophysin was measured within single pre-synapses (**F, L**). N=3, n=9 and 9 (48h), 9 and 6 (8d), for Ctr and CK1δ_i_, respectively. A nested t-test was used. **M** Quantification of tau+ neuron number upon CK1δ_i_ treatment for 6h, 24h, 48h or 8d analysed at day 15 normalised to untreated per timepoint. N=3 and n=8 and 8 (6h), 8 and 8 (24h), 8 and 7 (48h), 8 and 8 (8d), for Ctr and CK1δ_i_, respectively. A nested t-test was used. **N** Schematic of the timeline of CK1δ_i_. **O, P** Quantification of the percentage of pPERK+ GVB+ neurons (**O**) and mean nuclear CK1δ intensity (**P**) upon CK1δ_i_ treatment analysed at day 22 normalised to untreated per timepoint. N=3 and n=9. A nested t-test was used. Data are presented as mean ± SEM. *p*-values are indicated: \**p* < 0.05, \*\*\**p* < 0.001, \*\*\*\**p* < 0.0001 and ns: not significant.

**Figure S3:**
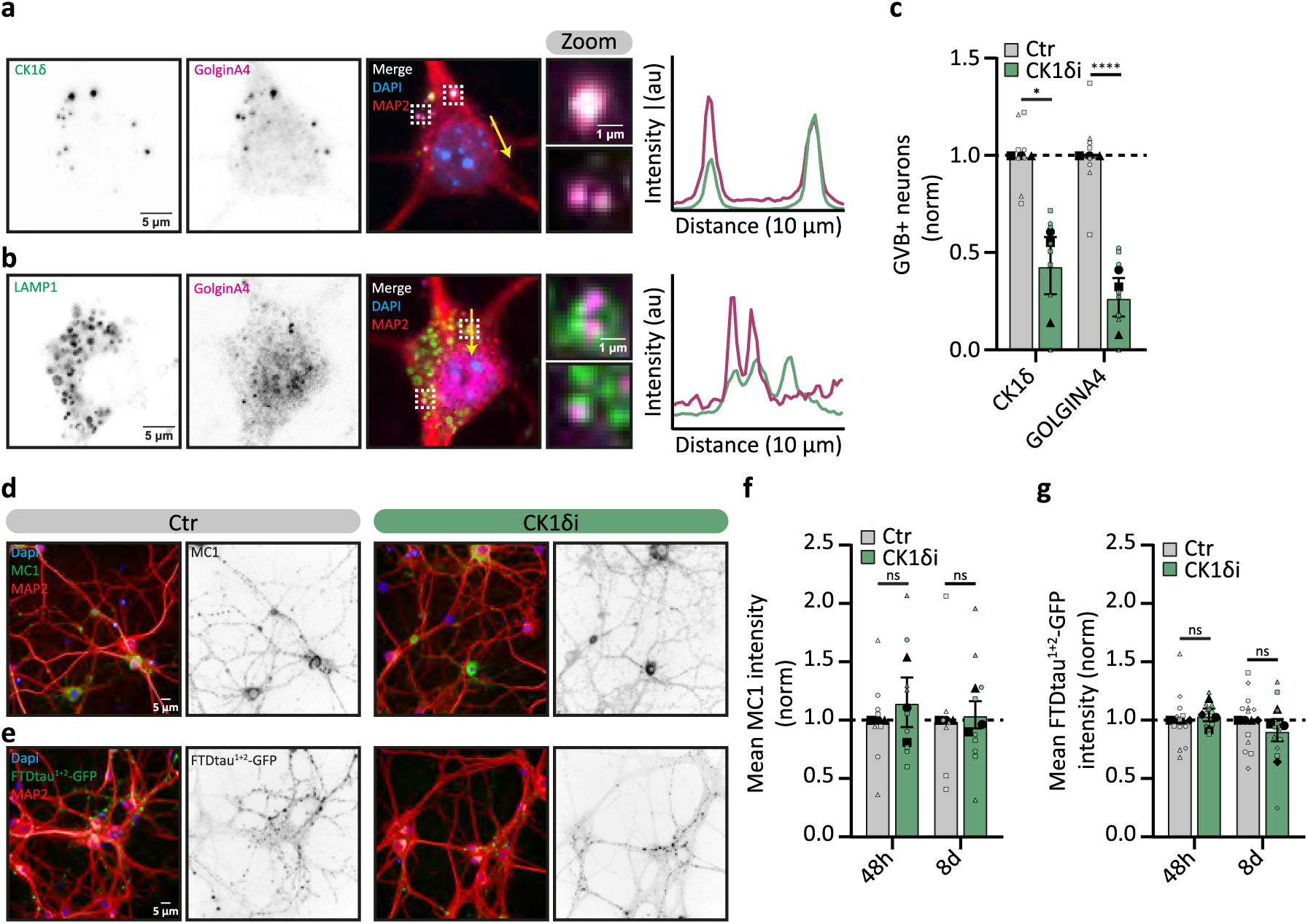
CK1δ inhibition does not affect GVB detection or tau pathology, related to Figure 2. **A, B** Representative confocal images of tau+/GVB+ neurons analysed on day 22. Neurons were immunostained for the GVB markers CK1δ (green) and GOLGINA4 (magenta) (**A**), the lysosomal membrane marker LAMP1 (green) and GOLGINA4 (magenta) (**B**) and the dendrite marker MAP2 (red). The zoomed regions are indicated by dashed white squares and the yellow arrow represents the location of the line intensity profiles. **C** Quantification of the percentage of CK1δ+ or GOLGINA4+ GVB+ neurons upon 48h of CK1δ_i_ treatment analysed at day 22 normalised to untreated per GVB-marker. N=3, n=8. A nested t-test was used. **D, E** Representative high-content microscopy images of FTDtau^1+2^- (**D**) or FTDtau^1+2^-GFP- (**E**) transduced (green) neurons treated without or with CK1δ_i_ for 48h and analysed at day 15. Neurons were immunostained for the pathological tau marker MC1 (green) (**d**) and MAP2 (red). **F** Quantification of mean MC1 immunofluorescence intensity upon CK1δ_i_ for 48h or 8d normalised to untreated control. N=3 and n=8 and 9 (48h), 7 and 9 (8d), for Ctr and CK1δ_i_, respectively. A nested t-test was used. **G** Quantification of mean FTDtau^1+2^-GFP intensity upon CK1δ_i_ for 48h or 8d normalised to untreated control. Single data points represent one well. N=4 and n=12 and 10 (48h), 12 and 11 (8d), for Ctr and CK1δ_i_, respectively. A nested t-test was used. Nuclei were visualised by DAPI (blue) and separate channels are shown in greyscale (**A, B, D, E**). Data are presented as mean ± SEM. *p*-values are indicated: \**p* < 0.05, \*\**p* < 0.01 and ns: not significant.

**Figure S4:**
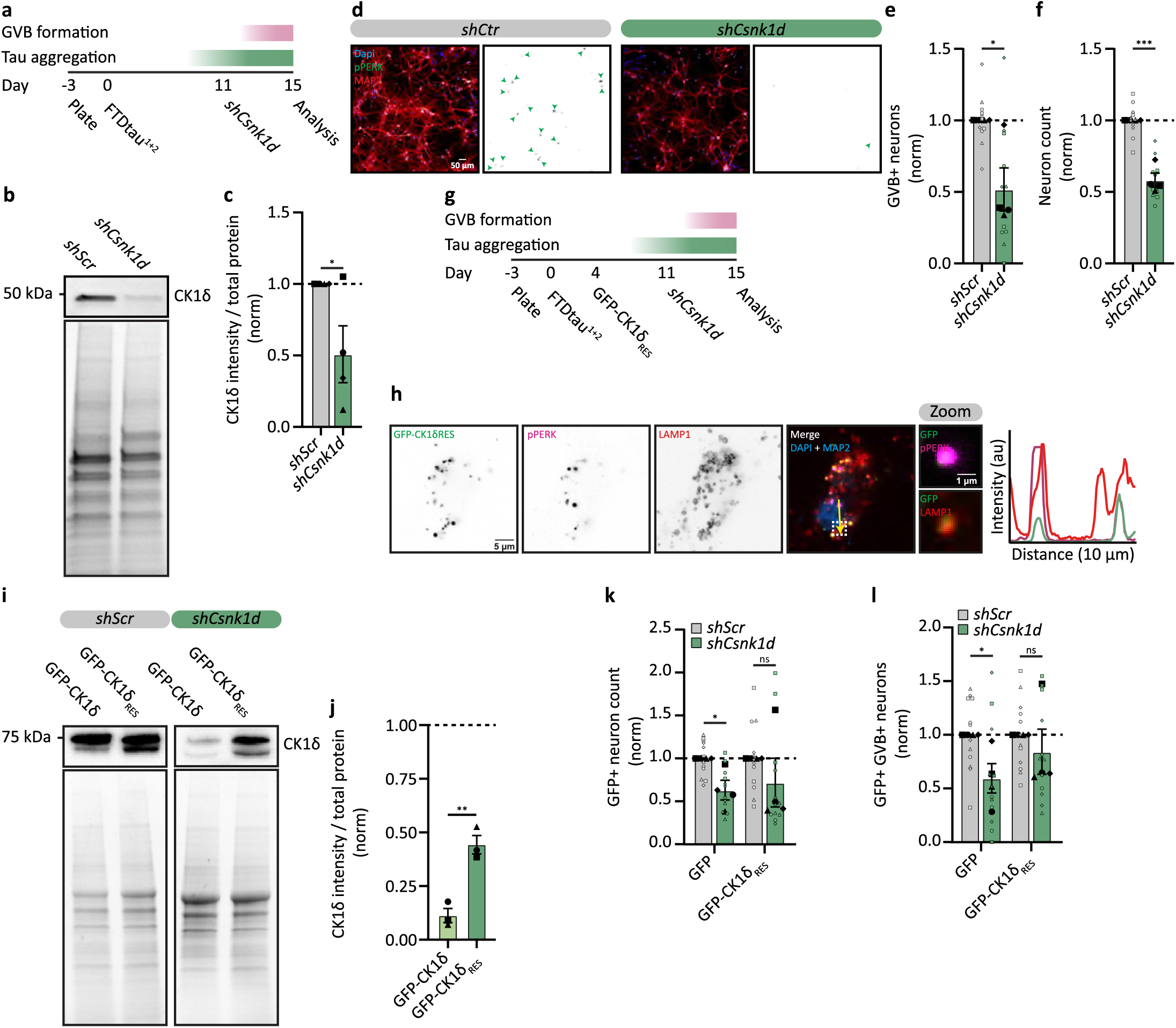
CK1δ knock-down reduces GVB formation, related to Figure 2. **A** Schematic of the timeline of CK1δ knock-down (KD). **B** Representative western blot of CK1δ of neurons transduced with *shScr* or *shCsnk1d* 4d before analysis at day 15. **C** Quantification of CK1δ intensity over total protein upon *Csnk1d* KD normalised to *shScr*. N=4. An unpaired t-test was used. **D** Representative high-content microscopy images of tau+ neurons transduced with *shScr* or *shCsnk1d* 4d before analysis at day 15. Neurons were immunostained for the GVB marker pPERK (green) and the dendrite marker MAP2 (red). Green arrowheads indicate GVB+ neurons. **E, F** Quantification of the percentage of pPERK+ GVB+ neurons (**E**) or neuron number (**G**) of tau+ neurons upon 4d of *Csnk1d* KD normalised to *shScr*. Single datapoints represent one well. N=4 and n=11 and 12, for *shScr* and *shCsnk1d*, respectively. A nested t-test was used. **G** Schematic of the timeline of CK1δ KD and rescue. **H** Representative confocal images of tau+/GVB+ neurons analysed at day 22. Neurons were transduced with GFP-CK1δ_RES_ which also localised to GVBs. The neurons were immunostained for pPERK (magenta), the lysosomal membrane marker LAMP1 (red) and MAP2 (blue). The zoomed region is indicated by a dashed white square and the yellow arrow represents the location of the line intensity profile. **I** Representative western blot of CK1δ of neurons transduced with GFP-CK1δ or GFP-CK1δ_RES_ at day 4 and subsequently transduced with either *shScr* or *shCsnk1d* at day 11 and analysed at day 15. **J** Quantification of CK1δ intensity over total protein upon *Csnk1d* KD normalised to *shScr* per group and analysed at day 15. GFP-CK1δ_RES_ shows a partial rescue of CK1δ protein levels. N=3. An unpaired t-test was used. **K, L** Quantification of the number of GFP+ neurons (**K**) or the percentage of pPERK+ GVB+ neurons (**L**) transduced with GFP or GFP-CK1δ_RES_ upon *Csnk1d* KD analysed at day 15 and normalised to *shScr* transduced neurons per group. N=4 and n=12. Nuclei were visualised by DAPI (blue) and separate channels are shown in greyscale (**D, H**). Data are presented as mean ± SEM. *p*-values are indicated: \**p* < 0.05, \*\**p* < 0.01, \*\*\**p* < 0.001 and ns: not significant.

**Figure S5:**
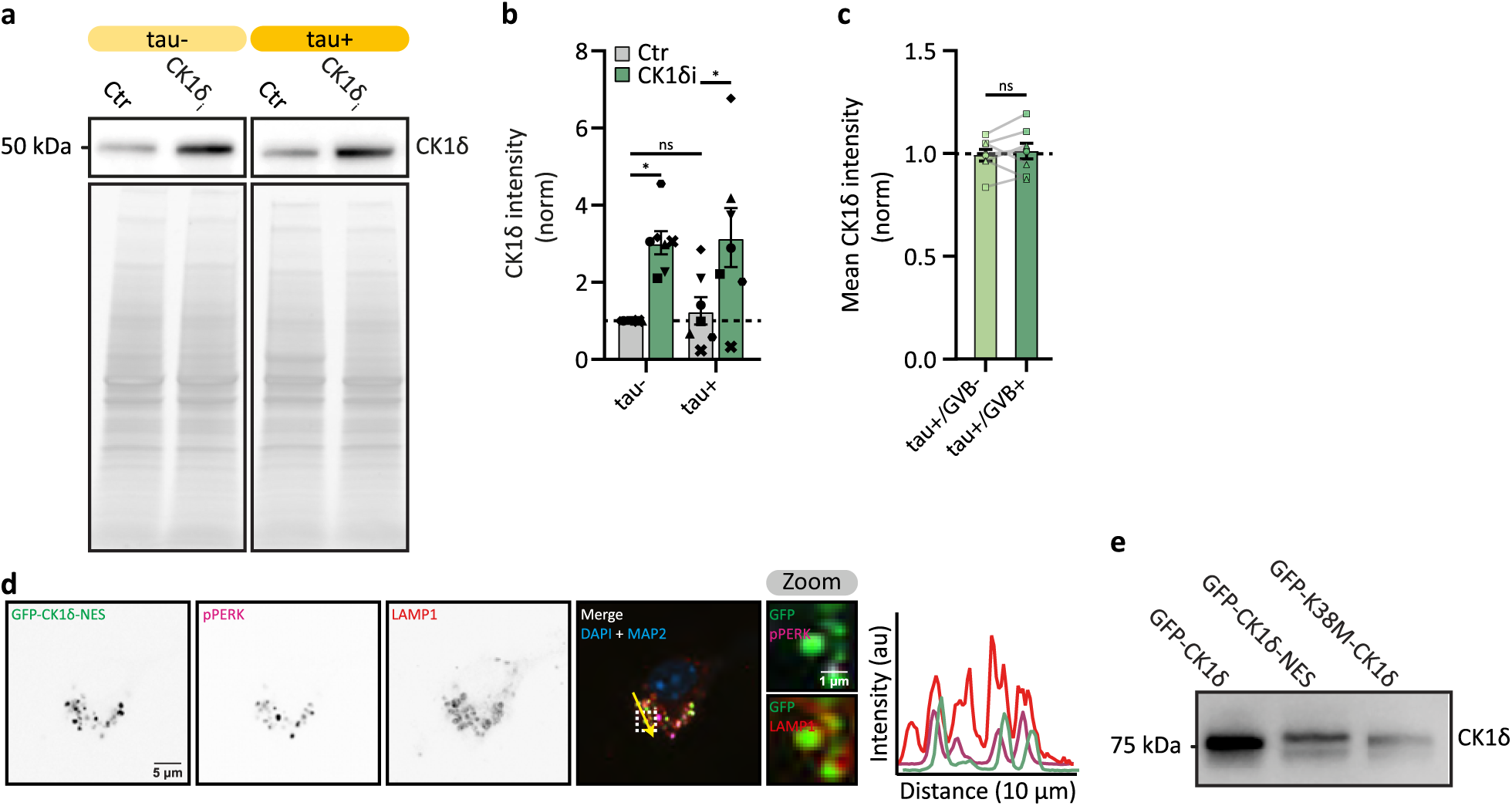
Tau aggregation does not increase CK1δ levels, related to Figure 2. **A** Representative western blot of CK1δ of tau- or tau+ neurons that were treated without or with CK1δ_i_ for 48h. **B** Quantification of CK1δ intensity of tau- or tau+ neurons that were treated without or with CK1δ_i_ for 48h and analysed at day 15. N=7. A one-way ANOVA followed by a Sidak’s post-hoc test was used. **C** Quantification of somatic CK1δ intensity in tau+/GVB- and tau+/GVB+ (pPERK based) neurons, excluding the GVB-area, normalised to tau+/GVB-neurons. N=3 and n=8. A paired t-test was used. **D** Representative confocal images of tau+/GVB+ neurons analysed at day 22. Neurons were transduced with GFP-CK1δ-NES. The neurons were immunostained for the GVB marker pPERK (magenta), the lysosomal membrane marker LAMP1 (red) and the dendrite marker MAP2 (blue). Nuclei were visualised by DAPI (blue). Separate channels are shown in greyscale. The zoomed regions are indicated by dashed white squares and the yellow arrow represents the location of the line intensity profiles. **E** Representative western blots of CK1δ of neurons transduced with GFP-CK1δ, GFP-CK1δ-NES or GFP-CK1δ-K38M and analysed at day 15. Data are presented as mean ± SEM. *p*-values are indicated: \**p* < 0.05 and ns: not significant.

**Figure S6:**
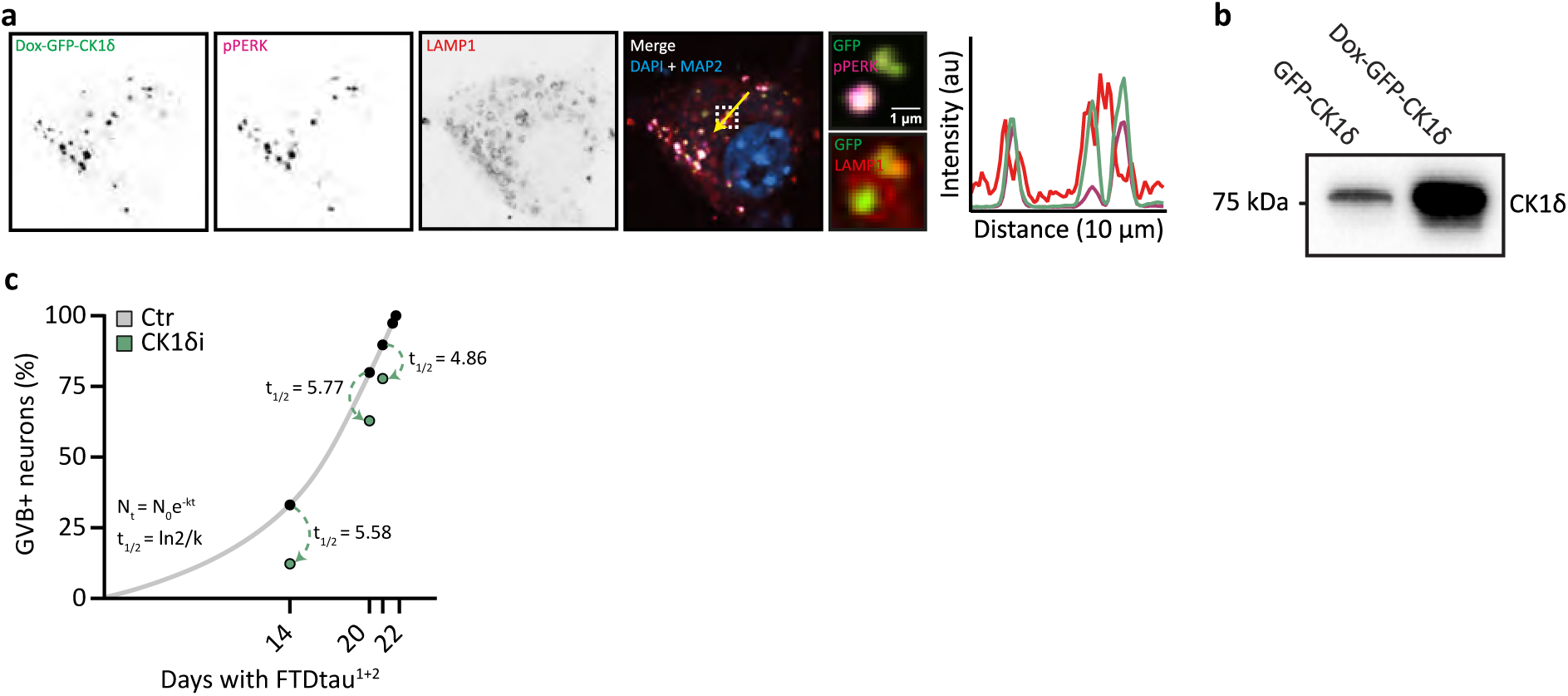
GVBs are stable structures, related to Figure 3. **A, B** Representative confocal images of tau+/GVB+ neurons analysed at day 22. Neurons were transduced with Dox-GFP-CK1δ and treated with Dox for 48h. The neurons were immunostained for the GVB marker pPERK (magenta), the lysosomal membrane marker LAMP1 (red) and the dendrite marker MAP2 (blue). Nuclei were visualised by DAPI (blue). Separate channels are shown in greyscale. The zoomed regions are indicated by dashed white squares and the yellow arrow represents the location of the line intensity profiles. **B** Representative western blots of CK1δ of neurons transduced with Dox-GFP-CK1δ and treated with Dox for 48h and analysed at day 15. **C** Visualisation of GVB half-life (t_1/2_) based on the polynomial equation obtained from Figure 1H (y = 0.05x^2^ - 0.3x + 0.2) to calculate the expected amount of GVB+ neurons after different exposure periods to tau aggregation (grey line) and the observed amount of GVB+ neurons upon 24h, 48h and 8d of CK1δ_i_ (Figure S2O). Assuming an exponential decay (N_t_=N_0_e^-kt^ and t_1/2_=ln2/k), t_1/2_ was calculated using the ratio of expected over observed amount of GVB+ neurons upon CK1δ_i_. The dotted green arrows indicate the difference in expected and observed amount of GVB+ neurons per timepoint.

**Figure S7:**
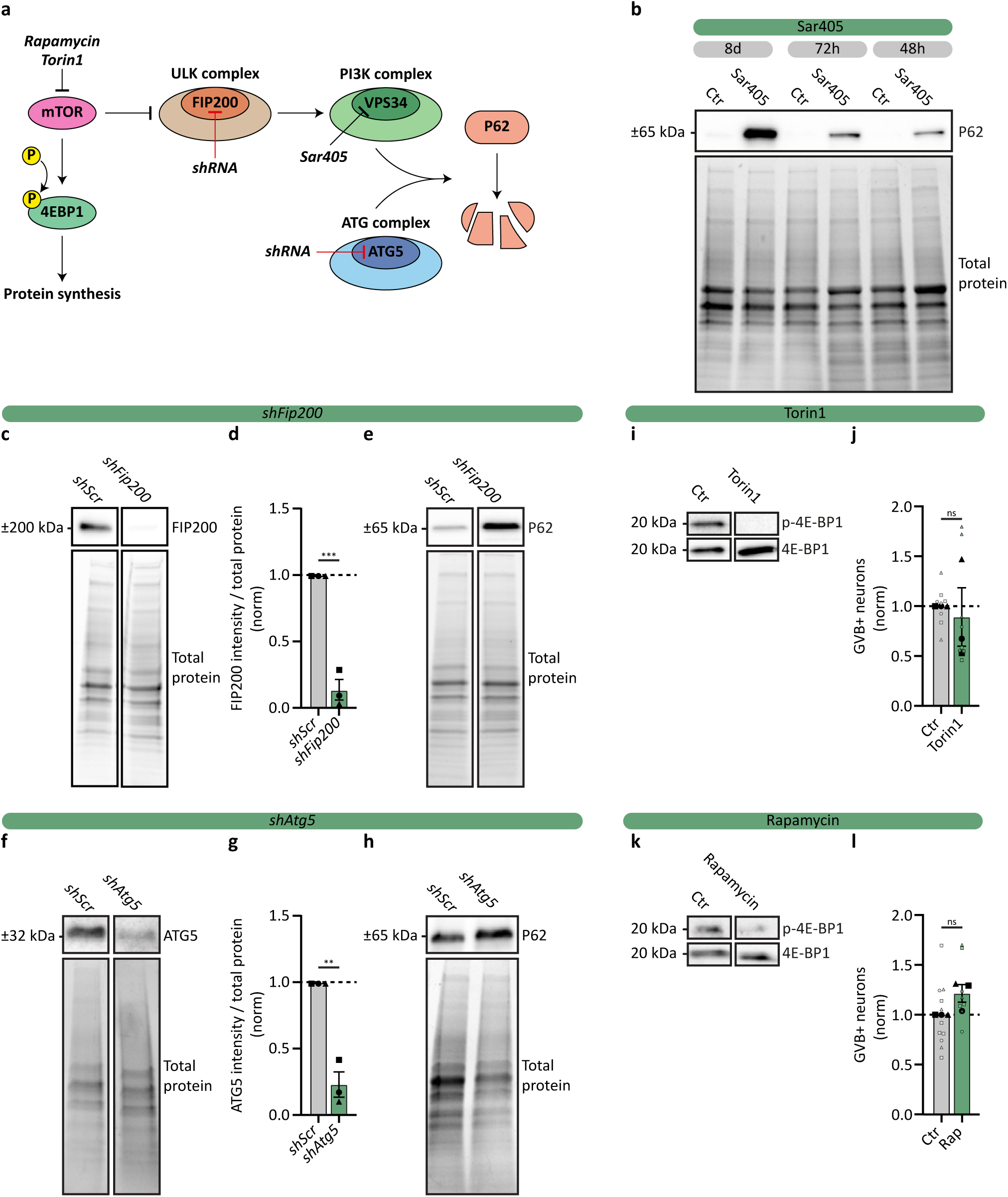
Validation of autophagy interventions, related to Figure 4. **A** Schematic of interventions used to block mTOR-dependent autophagy. **B** Representative western blot of P62 of neurons treated without or with Sar405 for 8d, 72h or 48h, analysed at day 15. **C** Representative western blot of FIP200 of neurons transduced with *shScr* or *shFip200* 8d before analysis at day 15. **D** Quantification of FIP200 intensity over total protein upon *Fip200* knock-down (KD) normalised to *shScr*. N=3. An unpaired t-test was used. **E** Representative western blot of P62 of neurons transduced with *shScr* or *shFip200* 8d before analysis at day 15. **F** Representative western blot of ATG5 of neurons transduced with *shScr* or *shAtg5* 8d before analysis at day 15. **G** Quantification of ATG5 intensity over total protein upon *Atg5* KD normalised to *shScr*. N=3. An unpaired t-test was used. **H** Representative western blot of P62 of neurons transduced with *shScr* or *shAtg5* 8d before analysis at day 15. **I, K** Representative western blot of p-4E-BP1 and 4E-BP1 of neurons treated without or with Torin1 (**I**) or rapamycin (**K**) for 48h analysed at day 15. **J, L** Quantification of the percentage of pPERK+ (**J**) or CK1δ+ (**L**) GVB+ neurons upon 48h of Torin1 (**J**) or rapamycin (**L**) treatment normalised to untreated per timepoint analysed at day 15.. N=3 and n=8 and 8 (**J**) or n=11 and 8 (**L**), Ctr and Torin1 or rapamycin, respectively. A nested t-test was used. Data are presented as mean ± SEM. *p*-values are indicated: \*\**p* < 0.01, \*\*\**p* < 0.001 and ns: not significant.

**Figure S8:**
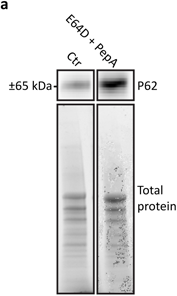
lysosomal inhibition using E64D and Pepstatin A reduces P62 degradation, related to Figure 4. **A** Representative western blot of P62 of neurons treated without or with E64D and Pepstatin A (PepA) for 24h, analysed at day 15.

**Figure S9:**
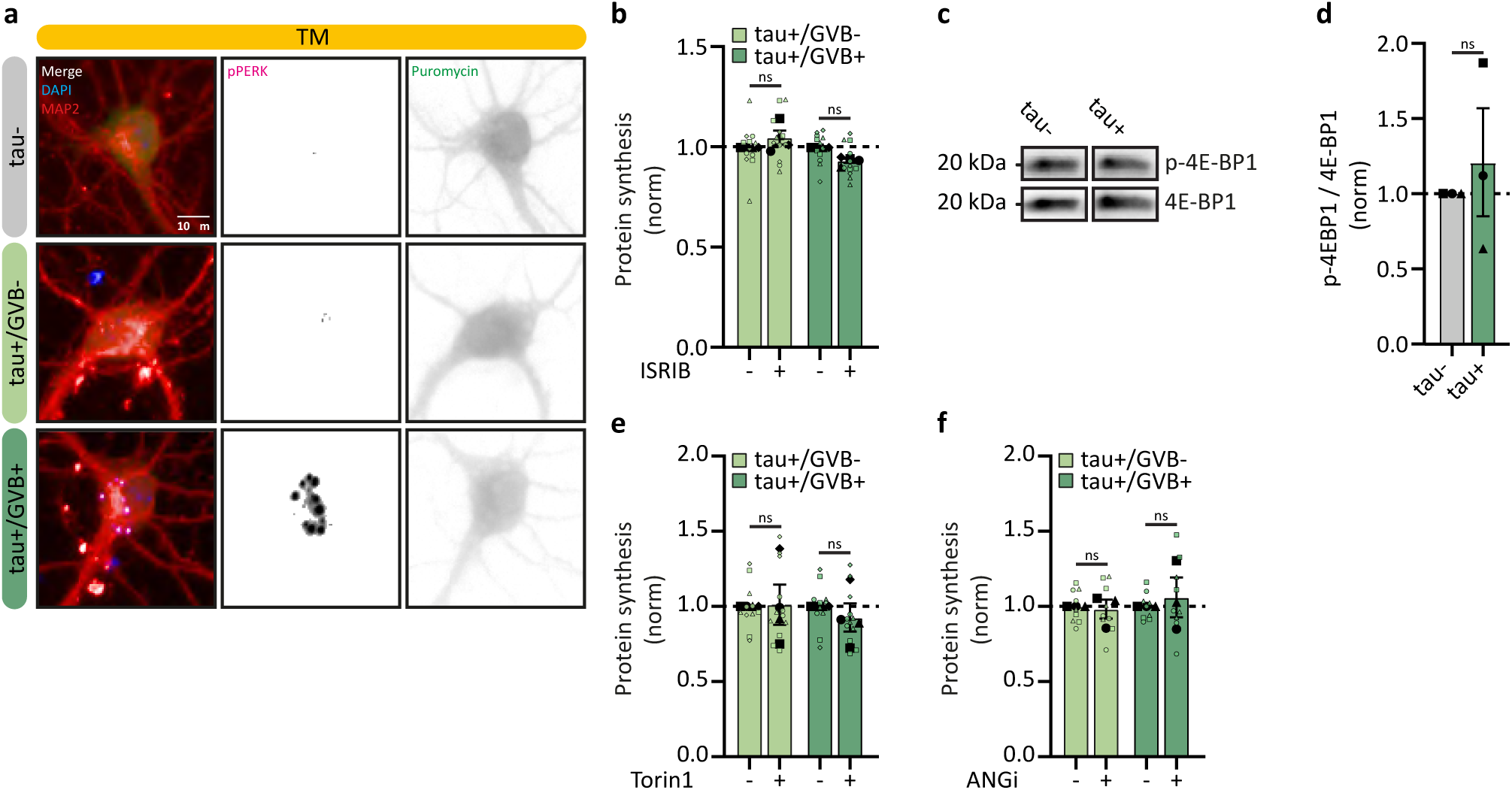
GVB formation and regulation does not depend on transient stress pathways, related to Figure 5. **A** Representative high-content microscopy images showing tau-, tau+/GVB- and tau+/GVB+ neurons treated with TM for 24h analysed at day 22. Neurons were treated for 15min with puromycin and were immunostained for puromycinilated proteins (green), the dendrite marker MAP2 (red) and the GVB marker pPERK (magenta). Nuclei were visualised by DAPI (blue). Separate channels are shown in greyscales. **B** Quantification of puromycin intensity as a measure for *de novo* protein synthesis in tau+/GVB- and tau+/GVB+ (pPERK based) neurons treated with the ISR inhibitor ISRIB for 2h analysed at day 22, normalised to untreated per group. N=3 and n=9. A nested t-test was used. **C** Representative western blot of p-4E-BP1 and 4E-BP1 of tau- or tau+ neurons analysed at day 15. **D** Quantification of p-4E-BP1 over 4E-BP1 intensity of tau+ neurons normalised to tau-analysed at day 15. N=3. An unpaired t-test was used. **E, F** Quantification of puromycin intensity as a measure for *de novo* protein synthesis in tau+/GVB- and tau+/GVB+ (pPERK based) neurons treated with angiogenin inhibitor (ANG_i_, **E**) or Torin1 (**F**) for 24h analysed at day 22, normalised to untreated per group. N=3 and n=9. A nested t-test was used. Data are presented as mean ± SEM. *p*-values are indicated: ns: not significant.

**Figure S10:**
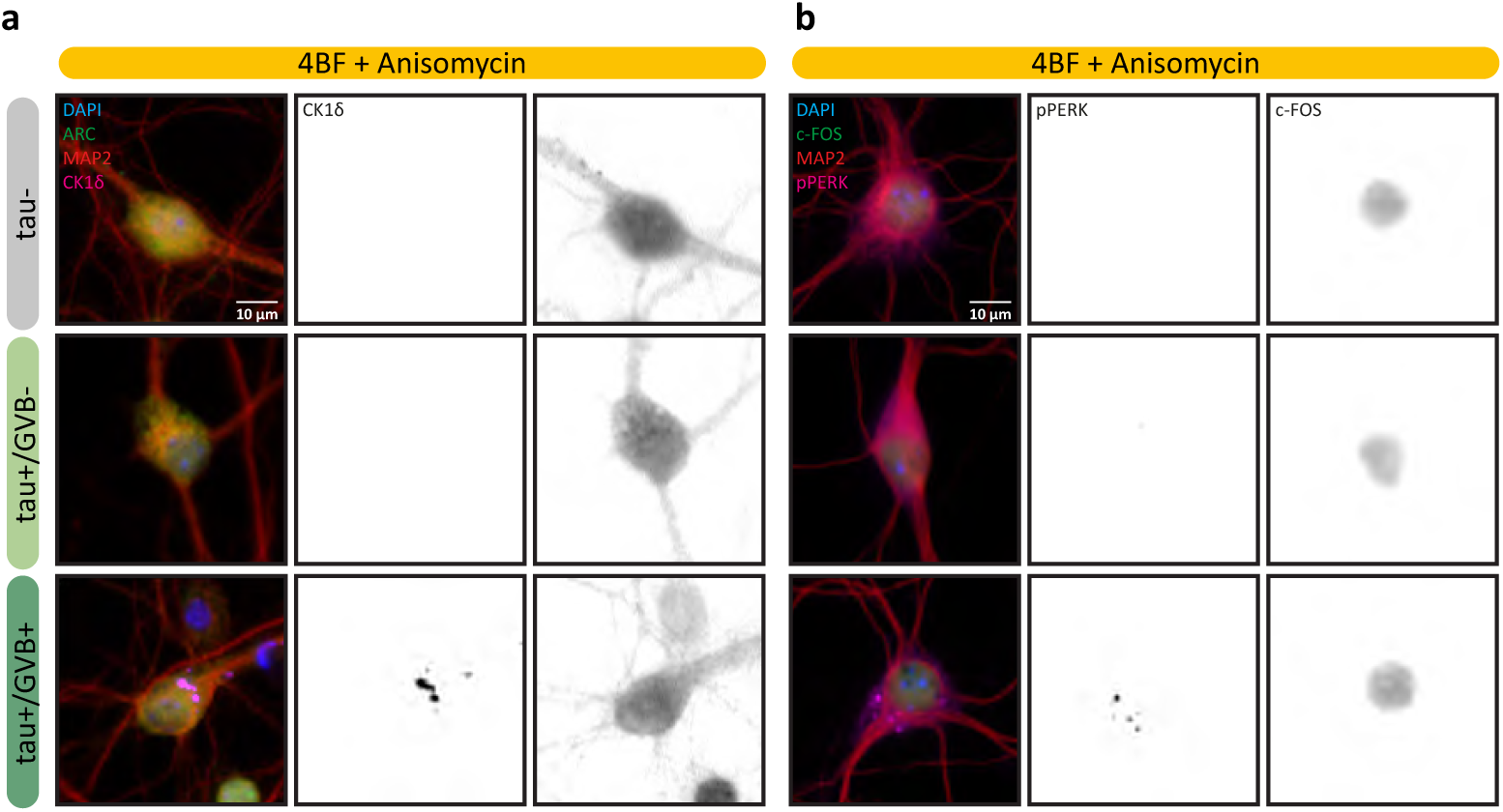
LTP-dependent induction of immediately early genes (IEGs) is dependent on protein synthesis, related to Figure 6. **A, B** Representative high-content microscopy images of tau-, tau+/GVB- and tau+/GVB+ neurons treated with anisomycin prior to treatment with 4BF (see Methods) for 4h. Neurons were immunostained for IEGs ARC (**A**) or c-FOS (**B**) (green), the dendrite marker MAP2 (red) and the GVB markers CK1δ (magenta) (**A**) or pPERK (magenta) (**A**). Nuclei were visualised by DAPI (blue). Separate channels are shown in greyscales.

**Video S1: GVBs are stable, dynamic structures, related to Figure 3**

Representative live imaging of a tau+ neuron transduced with GFP-CK1δ (grey) and imaged at day 21. An image was taken every 20 min for the duration of 9.20h. Although GVBs very rarely appeared or disappeared, this example shows a GVB disappearing (red arrow) and one appearing (black arrow).

